# A peptide inhibitor of Tau-SH3 interactions ameliorates amyloid-β toxicity

**DOI:** 10.1101/825760

**Authors:** Travis Rush, Jonathan R. Roth, Samantha J. Thompson, Adam R. Aldaher, J. Nicholas Cochran, Erik D. Roberson

## Abstract

The microtubule-associated protein Tau is strongly implicated in Alzheimer’s disease (AD) and aggregates into neurofibrillary tangles in AD. Genetic reduction of Tau is protective in several animal models of AD and cell culture models of amyloid-β (Aβ) toxicity, making it an exciting therapeutic target for treating AD. A variety of evidence indicates that Tau’s interactions with Fyn kinase and other SH3 domain–containing proteins, which bind to PxxP motifs in Tau’s proline-rich domain, may contribute to AD deficits and Aβ toxicity. Thus, we sought to determine if inhibiting Tau-SH3 interactions ameliorates Aβ toxicity. We developed a peptide inhibitor of Tau-SH3 interactions and a proximity ligation assay (PLA)-based target engagement assay. Then, we used membrane trafficking and neurite degeneration assays to determine if inhibiting Tau-SH3 interactions ameliorated Aβ oligomer (Aβo)-induced toxicity in primary hippocampal neurons from rats. We verified that Tau reduction ameliorated Aβo toxicity in neurons. Using PLA, we identified a peptide inhibitor that reduced Tau-SH3 interactions in HEK-293 cells and primary neurons. This peptide reduced Tau phosphorylation by Fyn without affecting Fyn’s kinase activity state. In primary neurons, endogenous Tau-Fyn interaction was present primarily in neurites and was reduced by the peptide inhibitor, from which we inferred target engagement. Reducing Tau-SH3 interactions in neurons ameliorated Aβo toxicity by multiple outcome measures, namely Aβo-induced membrane trafficking abnormalities and neurite degeneration. Our results indicate that Tau-SH3 interactions are critical for Aβo toxicity and that inhibiting them is a promising therapeutic target for AD.

---

The microtubule-associated protein Tau canonically stabilizes microtubules (1) and regulates many cellular processes beyond microtubule dynamics including axonal transport, DNA maintenance, and both pre- and post-synaptic signaling (2). Mutations in Tau lead to the neurodegenerative disease frontotemporal dementia (FTD) (3) and Tau inclusions are found in many neurodegenerative disorders including FTD (4), Alzheimer’s disease (AD) (5, 6), and temporal lobe epilepsy (7). Therapies for these diseases are desperately needed and while Tau is an exciting potential target, the question of how to best target Tau remains open.

The discovery that genetic reduction of Tau prevents behavioral and cognitive deficits in an AD mouse model provided an important insight with translational implications (8). This finding has been replicated in multiple AD models (9-14) as well as in other disease models including epilepsy (15, 16). Tau reduction also ameliorates network hyperexcitability, a potential early driver of AD deficits (17), in AD models (8, 11, 12) and in wild-type mice (18, 19). Thus, Tau’s role in neurodegeneration may be due to its influence on network hyperexcitability.

Due to the beneficial effects of Tau reduction across diverse models, many studies have examined its underlying mechanisms, seeking to replicate its beneficial effects therapeutically. However, efforts targeting Tau post-translational modifications, Tau aggregation, or microtubule stabilization have not yet been clinically successful (20), highlighting the importance of developing new approaches to target Tau therapeutically (21).

One such approach is inhibiting the interaction between Tau and other AD-related proteins. Tau interacts with several proteins implicated in AD, many of which contain SH3 domains and bind Tau’s central proline-rich domain. The proline-rich domain is heavily post-translationally modified in AD (22) and has several PxxP motifs that mediate binding to SH3 domain-containing proteins, including Fyn kinase (12, 23, 24). Fyn is strongly implicated in AD and exogenous Aβo activates Fyn in dendritic spines, leading to NMDA receptor (NMDAR) phosphorylation and excitotoxicity in neurons (11, 12). Diverse evidence provides a strong premise for therapeutically targeting Tau-SH3 interactions, especially the Tau-Fyn interaction. Manipulating Tau or Fyn produces a converging phenotype: either Tau (25) or Fyn (26) knockout protects against Aβo toxicity, and both proteins promote network hyperexcitability (8, 11, 15, 27). Tau reduction prevents cognitive deficits caused by Fyn overexpression in an AD mouse model (11). In addition, Tau plays an important role in the dendrite and postsynapse (28) and mice expressing a truncated form of Tau that excludes Fyn from dendrites are protected against Aβ-induced cognitive deficits and seizure susceptibility (12), showing that these abnormalities may be influenced by Tau-SH3 interactions like that of Tau-Fyn. However, whether Tau-SH3 interactions directly contribute to these effects, as opposed to more indirect mechanisms, remains unknown.

In this study, we sought to determine the effects of inhibiting Tau-SH3 interactions, a novel target for treating AD. Since Tau reduction ameliorates Aβo toxicity in primary neurons (25), we used that system to explore a potential mechanism by which Tau reduction prevents Aβo toxicity. These studies provide insight to the mechanism by which Aβo toxicity is Tau-dependent and identify Tau-SH3 interactions as a potential therapeutic target for AD.

## Results

### Tau reduction ameliorates Aβo toxicity

We first replicated the beneficial effects of Tau reduction in a primary hippocampal neuron model of Aβo toxicity. We applied a previously published Tau antisense oligonucleotide (ASO) (18), which after 1 week of exposure reduced Tau levels to about half of normal levels (Figure 1A,B). Six days after treatment with either the Tau ASO or a nontargeting control (NTC) ASO (18), we applied 2.5 μM synthetic Aβo (characterized in Supplementary Figure 1) or their vehicle control for 24 hours. To measure Aβo toxicity, we used a modified MTT assay that provides a measure of membrane trafficking (30, 31). Briefly, neurons metabolize MTT into formazan, which can either remain in soluble intracellular vesicles or be exocytosed, forming insoluble crystals on the cell surface. Aβo increases exocytosis of formazan, thus decreasing the ratio of soluble vs insoluble formazan (see Supplementary Figure 2). Tau reduction ameliorated this Aβo-induced dysfunction, as the Tau ASO protected neurons from Aβo-induced membrane trafficking abnormalities (Figure 1C). Importantly, Tau reduction did not compromise viability of neurons, measured by either NeuN levels (Figure 1D) or MTT assay (Figure 1E). These results add to the body of work demonstrating that Tau reduction does not impair neuronal survival.

**Figure 1:**
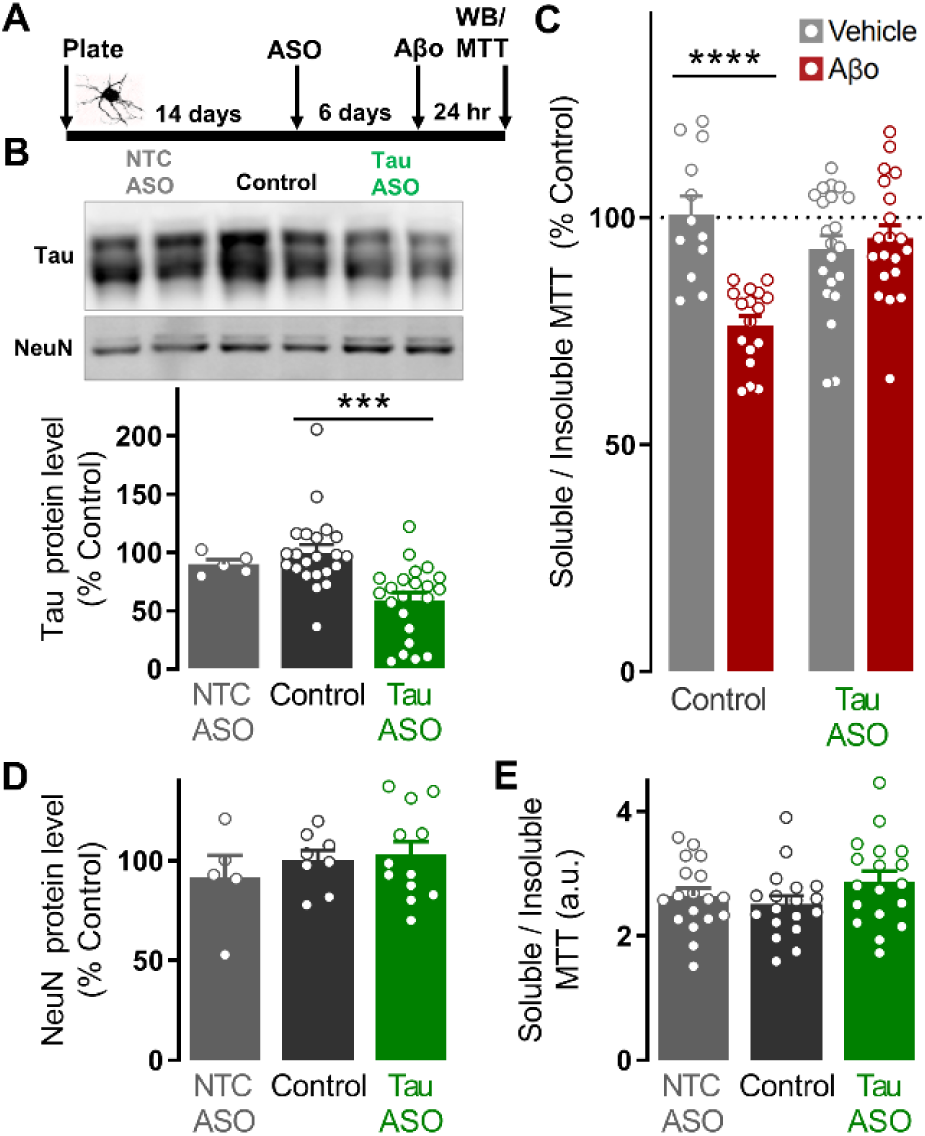
Tau reduction ameliorates amyloid-β–induced membrane trafficking abnormalities. **A:** Timeline of ASO treatment and modified MTT experiments. **B:** Representative Western blot images of lysate from hippocampal neurons from rats treated for one week (DIV14-21) with a previously published antisense oligonucleotide for Tau or nontargeting control ASO (18). The Tau ASO caused a 41% reduction in Tau while a nontargeting control (NTC) ASO did not reduce Tau (ANOVA, F(2,64) = 10.23, *p* = 0.0002, n = 5-22 wells per group). **C:** At DIV20, 2.5 μM Aβo or vehicle equivalent was applied. After 24 hours, Aβo-induced membrane trafficking abnormalities were measured using a modified MTT assay (full description in the methods section and Supplementary Figure 2). Tau reduction with the ASO ameliorated Aβo-induced membrane trafficking abnormalities (Two-way ANOVA interaction for factors Aβo and ASO F(1,66) = 18.9, *p* < 0.0001, n = 12-21 wells per group from N = 2 rats, 4-12 wells per group per plate with 1 plate per rat). Differences indicated: \*\*\*\**p* < 0.0001 by Dunnett’s *post hoc* test with correction for multiple comparisons relative to vehicle control. **D:** Neither ASO affected neuronal survival, as measured with NeuN protein levels (ANOVA, F(2,22) = 0.52, *p* = 0.60, n=5-12 wells per group). **E**: Neither ASO affected neuronal survival, as measured with the MTT assay (ANOVA, F(2,51) = 1.66, *p* = 0.42, n = 18 wells per group). All Panels: Bars indicate mean ± SEM. Differences indicated: \*\*\**p* < 0.001 by Dunnett’s *post hoc* test with correction for multiple comparisons relative to CTL.

### Tau-PxxP_5/6_ inhibits Tau-SH3 interactions

To determine whether the beneficial effects of Tau reduction would be replicated by inhibiting Tau-SH3 interactions, we developed a cell-permeable Tau-SH3 interaction inhibitor peptide (Tau-PxxP_5/6_) based on a 17–amino acid peptide spanning the 5^th^ and 6^th^ PxxP motifs of the proline-rich domain of human Tau protein (amino acids 209–225). This region of Tau is the primary binding site for Fyn and may mediate other Tau-SH3 interactions. We added a 10–amino acid sequence derived from the transactivator of transcription (TAT) of human immunodeficiency virus for cell permeability. We have previously shown that the resulting Tau-PxxP_5/6_ peptide inhibits Tau-Fyn interactions in a cell-free AlphaScreen (23). To determine whether Tau-PxxP_5/6_ inhibits the Tau-Fyn interactions in a cell-based system, we utilized proximity ligation assay (PLA) to measure the amount of Tau-Fyn interaction in cells. If two proteins interact and the employed antigen-targeting domains are within 40 nanometers of each other, PLA leads to a fluorescent punctum, which can be imaged by immunocytochemistry or immunohistochemistry. We transiently transfected HEK-293 cells with plasmids both for mKate2-tagged Tau and for Fyn (Figure 2A) and verified that the PLA signal was only seen in cells transfected with both Tau and Fyn and treated with primary antibodies (Supplementary Figure 3). Treatment of cells with Tau-PxxP_5/6_ for 24 hours decreased Tau-Fyn PLA signal by almost half (Figure 2B), measured as the number of PLA puncta per cell area. As a negative control to rule out a non-specific peptide effect, we also designed a cell-permeable TAT peptide (Tau-CTD) spanning 17 amino acids from Tau’s C-terminal domain (residues 380-396), a region that does not contain PxxP motifs or mediate Tau’s SH3 interactions. As expected, Tau-CTD did not change Tau-Fyn PLA density (Figure 2B). Tau-PxxP_5/6_ did not change Tau or Fyn levels in HEK-293 cells (Figure 2C), demonstrating that the PLA results are not due to reducing protein levels.

**Figure 2:**
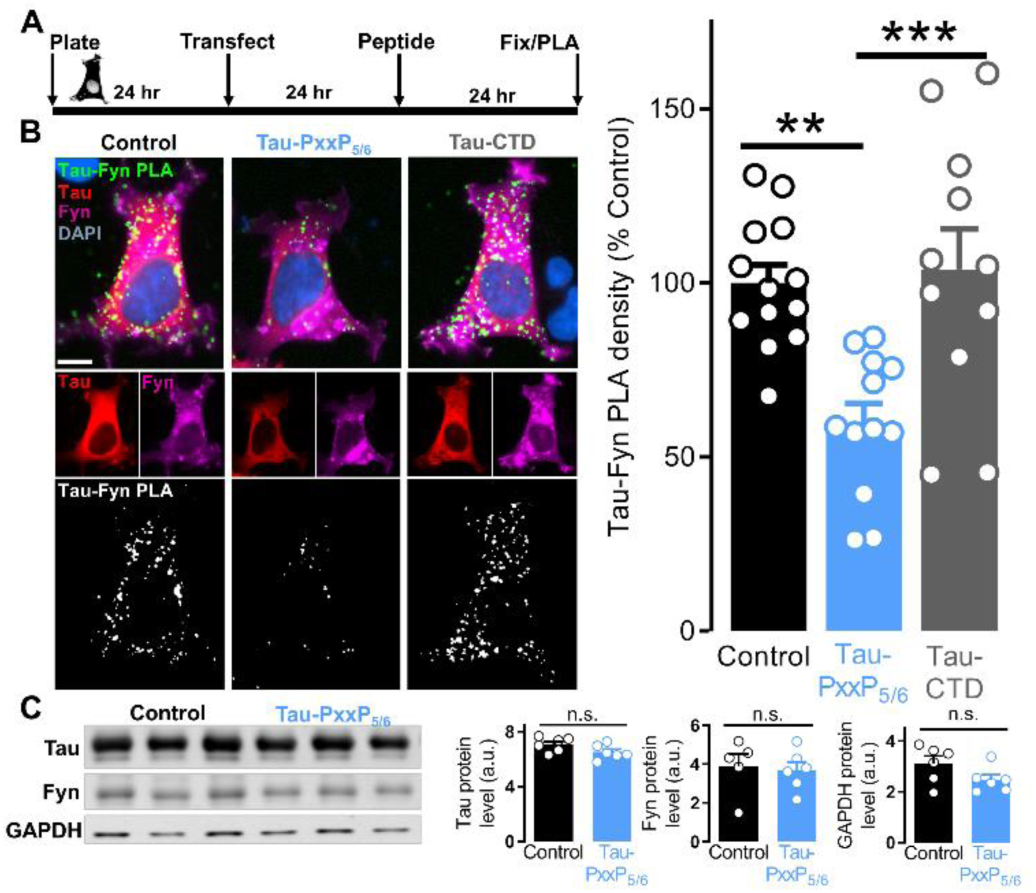
Tau-PxxP_5/6_ inhibits the Tau-Fyn interaction in cells. **A:** Timeline of Tau-Fyn PLA experiments in HEK-293 cells. **B:** Representative immunofluorescent images of HEK-293 cells treated with Tau-PxxP_5/6_ (rrrqrrkkrgRSRTPSLPTPPTREPKK) or Tau-CTD (rrrqrrkkrgENAKAKTD HGAEIVYKS) after Tau-Fyn proximity ligation assay (PLA) was performed (full description in the methods section). Green puncta represent sites of Tau-Fyn interaction, red is Tau-mKate2, and magenta is immunofluorescently labeled Fyn. Scale bar = 10 μm. Quantification of Tau-Fyn PLA density demonstrates that Tau-SH3 inhibitor Tau-PxxP_5/6_ reduced Tau-Fyn interaction by 40% while negative control peptide Tau-CTD did not (ANOVA, F(2,33) = 10.14, *p =* 0.0004, average of 5-7 images per coverslip from n = 11-13 coverslips from N = 6 distinct passages, see Methods for details of quantification and analysis). Differences indicated: \*\**p* < 0.01, and \*\*\**p* < 0.001 by Dunnett’s *post hoc* test with correction for multiple comparisons relative to control. **C:** Western blot of lysate from HEK-293 cells demonstrates that Tau-PxxP_5/6_ did not change Tau, Fyn, or GAPDH levels (Student’s t-test, t(10)=1.86, *p* = 0.092; t(9)=0.28, *p* = 0.79; t(10) =1.78, *p* = 0.10, respectively, n=5-6 wells per group).

### Tau-PxxP_5/6_ reduces Tau phosphorylation by Fyn without changing Fyn kinase activity

Fyn phosphorylates Tau at Y18, and this phosphorylation has been implicated in NMDAR-dependent glutamate toxicity (32), so we determined whether Tau-PxxP_5/6_ reduced the phosphorylation of Tau Y18 by Fyn. It is likely that Tau-Fyn interactions facilitate subsequent Tau phosphorylation by Fyn, so we expected that inhibiting their interaction via the SH3 domain would decrease phosphorylation of Tau at Y18 by Fyn. To test this, we transiently transfected HEK-293 cells with Tau and Fyn, then determined the effect of 24 hour exposure to Tau-PxxP_5/6_ on tau phosphorylation measured by Western blot (Figure 3A). As expected, Tau-PxxP_5/6_ reduced Y18 phosphorylation of Tau by Fyn (Figure 3B).

**Figure 3:**
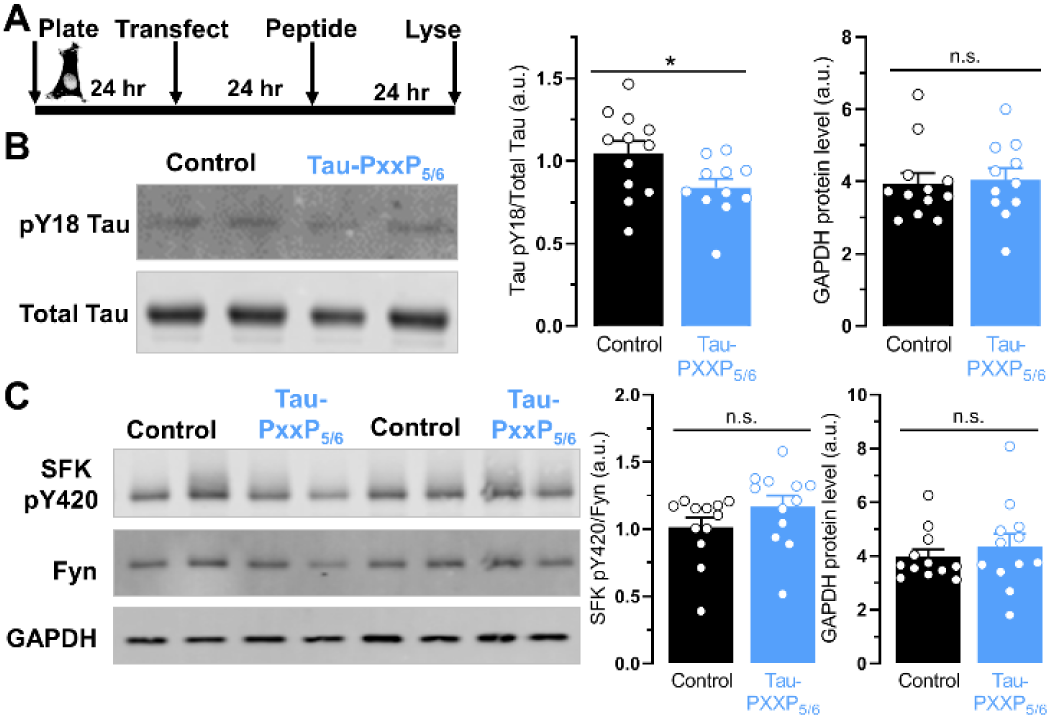
Tau-PxxP5/6 reduces phosphorylation of Tau by Fyn without changing Fyn kinase activity. **A:** Timeline of Tau and Fyn phosphorylation experiments in HEK-293 cells. **B:** Representative Western blot images of lysate from HEK-293 cells treated with Tau-PxxP_5/6_ and probed for Tau PY18 and total Tau. Tau-PxxP_5/6_ reduced phosphorylation of Tau pY18 by Fyn by 19.75% (Student’s t-test, t(21) = 2.23, *p* < 0.037, average of 2 technical replicates of n=11-12 wells per group) without affecting GAPDH protein level (Student’s t-test, t(21) = 0.25, *p* = 0.80, average of 2 technical replicates of n=11-12 wells per group). **C:** Representative Western blot images of lysate from HEK-293 cells treated with Tau-PxxP_5/6_ and probed for Src family kinase (SFK) pY420, Fyn, and GAPDH. Tau-PxxP_5/6_ did not affect Fyn pY420 (Student’s t-test, t(22) = 1.40, *p* = 0.17, n = 12 wells per group) or GAPDH (Student’s t-test, t(22) = 0.71, *p* = 0.48, n = 12 wells per group).

This effect likely results from reduced Tau-Fyn interaction but could be due to direct Fyn inhibition by Tau-PxxP_5/6_. To distinguish between these possibilities, we examined Fyn’s regulatory phosphorylation status. Fyn autophosphorylation at Y420 is associated with active Fyn kinase and assessing phosphorylation at this site is a commonly used measure of Fyn kinase activity (33). Importantly, we found that Tau-PxxP_5/6_ did not change phosphorylation status of Fyn at Y420, providing no evidence for a change in Fyn activity (Figure 3C). As positive controls, the Fyn kinase inhibitor PP2 decreased pY420 (Supplementary Figure 4), and ethanol, which activates Fyn kinase activity (34), increased pY420 (Supplementary Figure 5). Together, these results support the hypothesis that Tau-PxxP_5/6_ inhibits Tau-Fyn interaction without changing the activity of Fyn kinase.

### Tau-PxxP_5/6_ inhibits endogenous Tau-Fyn interaction in neurons

Since Tau-PxxP_5/6_ inhibits the Tau-Fyn interaction when the proteins are overexpressed in HEK-293 cells, we next used PLA to determine if Tau-PxxP_5/6_ inhibits endogenous Tau-Fyn interaction in neurons, achieving the intended pharmacodynamic effect and allowing us to infer target engagement. This also provided the opportunity to visualize the amount and localization of endogenous Tau-Fyn interaction in neurons (Figure 4A). Interestingly, the interaction predominantly occurred throughout the neurites rather than in the soma and appeared adjacent to, but not directly within, Tau-positive neuritic shafts (see inset). This localization is consistent with the possibility that the Tau-Fyn interaction may occur in dendritic spines, which would support the hypothesis that Tau and Fyn interact at the postsynapse in dendrites, allowing Aβo to transduce its signal through Fyn, leading to neuronal dysfunction.

**Figure 4:**
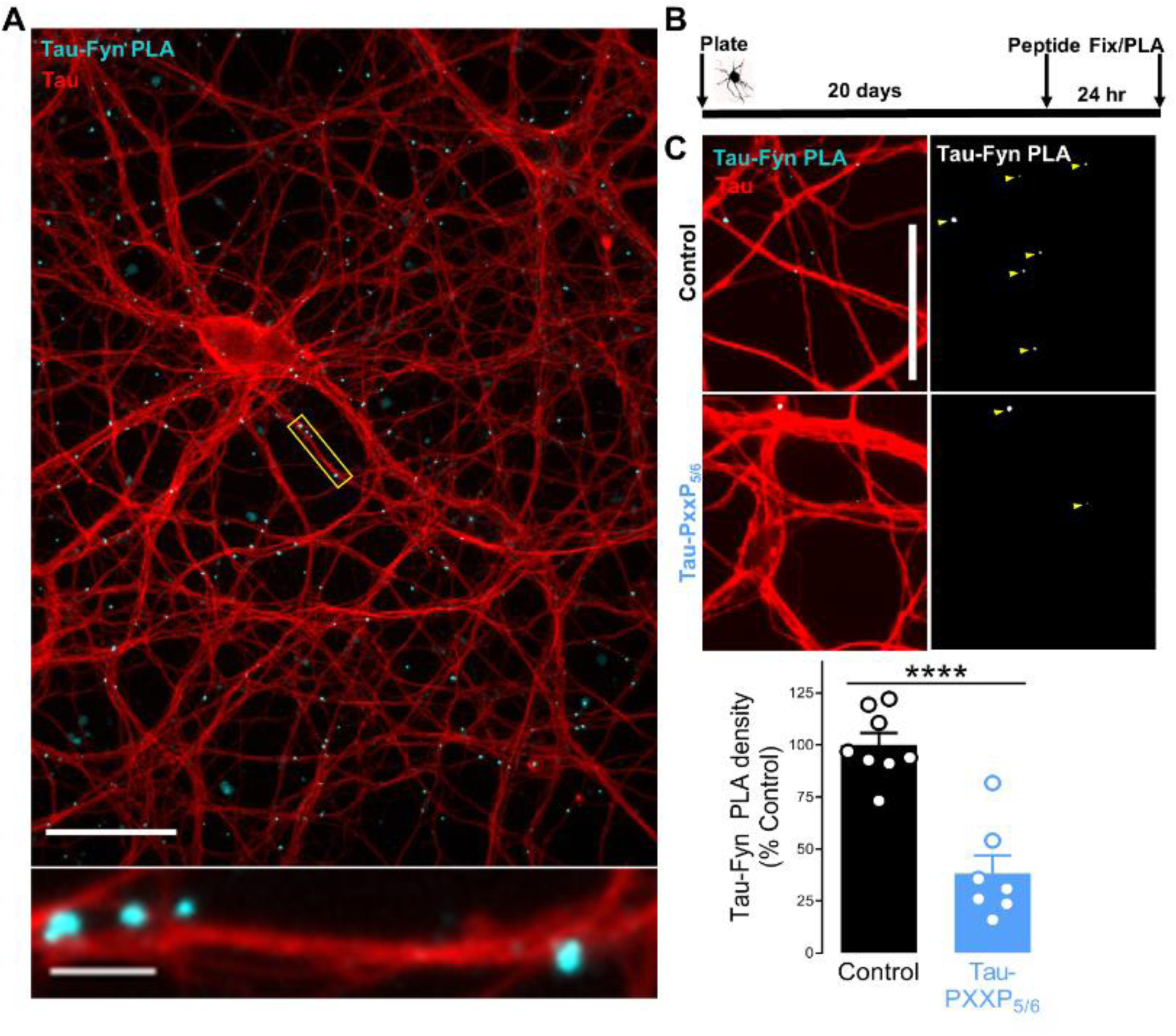
Tau-PxxP_5/6_ inhibits endogenous Tau-Fyn interaction in neurons. **A:** Representative immunofluorescent image of DIV21 primary neurons after Tau-Fyn PLA. Cyan puncta represent sites of Tau-Fyn interaction and red is immunofluorescently labeled Tau. Scale bar = 50 μm. *Inset:* magnification of a neurite showing Tau-Fyn PLA (cyan) adjacent to Tau-positive neurites (red). Scale bar = 5 μm. **B:** Timeline of endogenous Tau-Fyn PLA inhibition experiments. **C:** Representative images of endogenous Tau-Fyn PLA after Tau-PxxP_5/6_ treatment demonstrating reduction of endogenous Tau-Fyn PLA in neurons by Tau-PxxP_5/6_. Tau-PxxP_5/6_ reduced endogenous Tau-Fyn interaction in neurons by 62% (Student’s t-test, t(13) = 6.124, *p* < 0.0001, average of 5-7 images per coverslip from n = 7-8 coverslips). Scale bar = 25 μm.

Treating DIV20 neurons with Tau-PxxP_5/6_ for 24 hours decreased endogenous Tau-Fyn PLA density by about two-thirds (Figure 4B,C). Lower magnification images of neurons treated with Tau-PxxP_5/6_ demonstrate that the peptide did not affect the morphology of neurons (Supplemental Figure 6). Importantly, Tau-PxxP_5/6_ did not change Tau or Fyn levels in primary neurons (Supplementary Figure 7), and reducing Tau or Fyn with ASOs reduced Tau-Fyn PLA (Supplementary Figure 8). These results show that Tau and Fyn interact endogenously in neurons, that this interaction predominately occurs in neurites, and that Tau-PxxP_5/6_ inhibits endogenous Tau-Fyn interactions.

### Inhibiting Tau-SH3 interactions ameliorates Aβo toxicity

With validated tools in hand for inhibiting Tau-SH3 interactions in neurons, we next asked if inhibiting Tau-SH3 interactions ameliorated Aβo toxicity in primary neurons, mimicking the beneficial effects of Tau reduction. We assessed Aβo toxicity using two orthogonal measures: membrane trafficking abnormalities and neurite degeneration.

First, we tested whether inhibiting Tau-SH3 interactions prevented Aβo-induced membrane trafficking abnormalities. We pretreated DIV20 neurons with Tau-PxxP_5/6_ or its vehicle control for 90 minutes before applying 2.5 μM Aβo or their vehicle control for 24 hours and utilizing the same modified MTT assay described earlier (Figure 5A). Tau-PxxP_5/6_ ameliorated Aβo-induced membrane trafficking abnormalities in a dose-dependent manner, while the negative control peptide, Tau-CTD, did not (Figure 5B).

**Figure 5.**
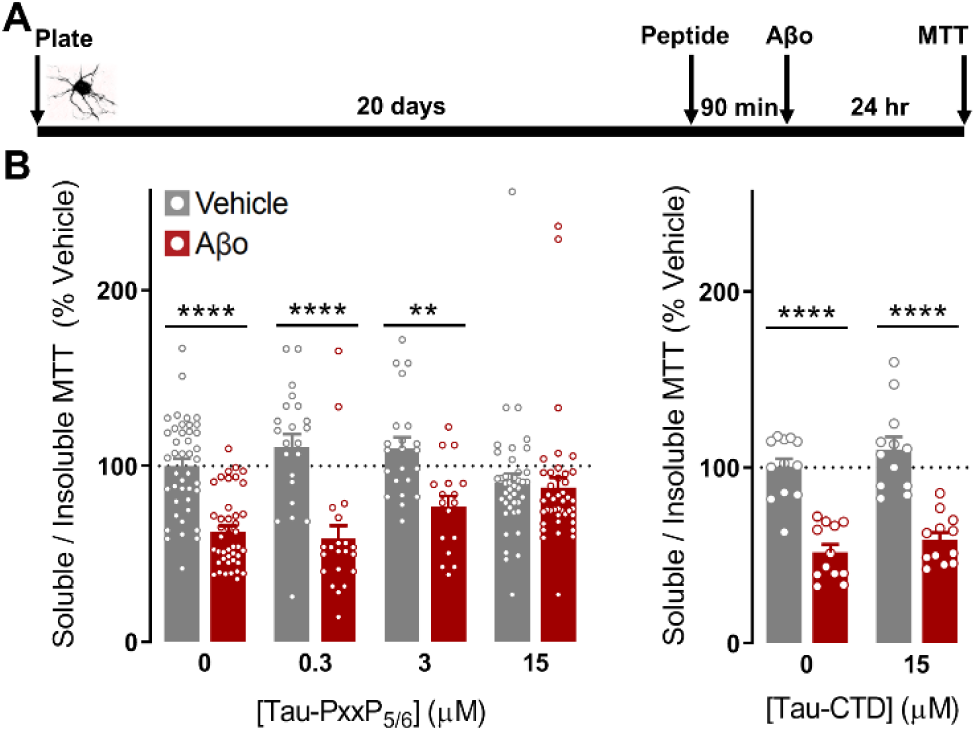
Inhibiting Tau-SH3 interactions ameliorates amyloid-β– induced membrane trafficking abnormalities. **A:** Timeline of Tau-PxxP_5/6_ modified MTT assay. Neurons were pretreated with increasing doses of Tau-PxxP_5/6_ for 90 minutes before 2.5 μM Aβo application. After 24 hours, the modified MTT assay detailed earlier was performed. **B:** Tau-PxxP_5/6_ prevented Aβo-induced membrane trafficking abnormalities in a dose-dependent fashion (Two-way ANOVA interaction for factors Aβo and Tau-PxxP_5/6_ F(3,241) = 7.74, *p* < 0.0001, n = 17-42 wells per group from N = 3 embryo harvests, 3-6 wells per group per plate with 2 plates per harvest for 2 harvests and 9-12 wells per group per plate with 2 plates per harvest for 1 harvest) while the negative control peptide Tau-CTD failed to ameliorate Aβo-induced membrane trafficking abnormalities (Two-way ANOVA interaction for factors Aβo and Tau-CTD F(1,44) = 0.092, *p* = 0.76, n = 12 wells per group from N=2 embryo harvests, 6 wells per group per plate with 1 plate per harvest). Differences indicated: \*\**p*<0.01, and \*\*\*\**p*<0.0001, by Dunnett’s *post hoc* test with correction for multiple comparisons relative to vehicle control at that dose of peptide. Bars indicate mean ± SEM

Second, we tested another Aβo-induced abnormality that is ameliorated by Tau reduction (25), Aβo-induced neurite degeneration. Aβo causes neurite degeneration and blebbing, which can be quantified using immunofluorescent labeling of MAP2 followed by imaging in an unbiased manner using a high-content, automated approach (see Supplementary Figure 9). As before, we pretreated DIV20 neurons with Tau-PxxP_5/6_ or its vehicle control for 90 minutes before applying 2.5 μM Aβo or their vehicle control (Figure 6A). Forty-eight hours after Aβo application, we fixed the neurons, labeled MAP2, then quantifiedfig intact neurite length. Tau-PxxP_5/6_ ameliorated neurite blebbing and degeneration while Tau-CTD did not (Figure 6B,C). Together, these findings provide evidence that Tau-SH3 interactions contribute to Aβo toxicity in neurons.

**Figure 6.**
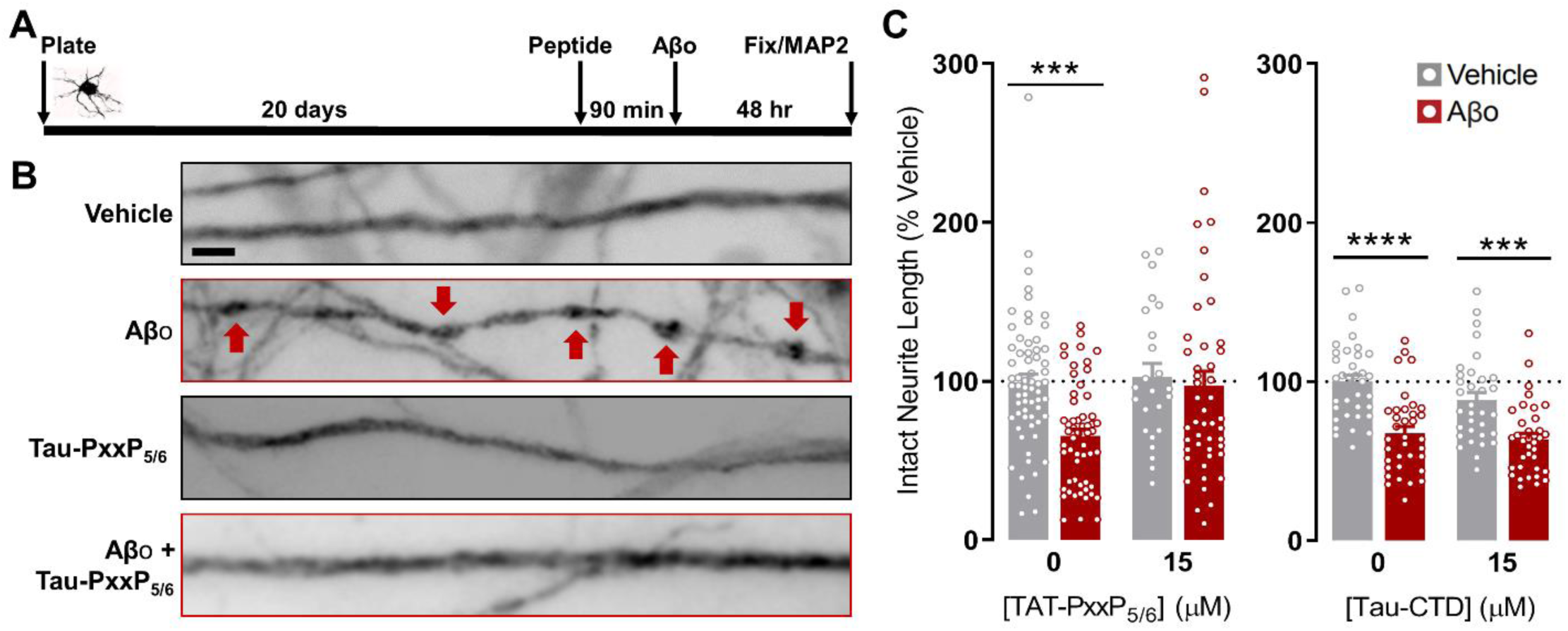
Inhibiting Tau-SH3 interactions ameliorates amyloid-β–induced neurite degeneration. **A:** Timeline of Tau-PxxP_5/6_ MAP2. **B:** Representative images of neurons show signs of neurite blebbing and degeneration by immunofluorescent labeling of MAP2 (grayscale, blebbing indicated by arrows) after 48 hour exposure to Aβo. Neurons pretreated with 15 μM Tau-PxxP_5/6_ showed no blebbing or signs of neurite degeneration. Scale bar = 5 μm. **C:** Unbiased quantification of intact neurite length from MAP2 staining (see Supplementary Figure 4) demonstrate that Tau-PxxP_5/6_ ameliorated Aβo-induced neurite degeneration (Two-way ANOVA interaction for factors Aβo and Tau-PxxP_5/6_ F(1,115) = 3.98, *p* = 0.048, n = 24-64 wells per group from N = 5 neuron harvests, 3-6 wells per group per plate with 2 plates per harvest for 4 harvests and 6-12 wells per group per plate with 2 plates per harvest for 1 harvest) while Tau-CTD did not (Two-way ANOVA interaction for factors Aβo and Tau-CTD F(1,135) = 0.857, *p* = 0.35, n = 33-36 wells per group from N = 3 embryo harvests, 11-12 wells per group per plate with 1 plate per harvest). Differences indicated: \*\*\**p*<0.001, and \*\*\*\**p*<0.0001, by Dunnett’s *post hoc* test with correction for multiple comparisons relative to vehicle control at that dose of peptide. Bars indicate mean ± SEM.

## Discussion

This study demonstrates that an inhibitor of Tau-SH3 interactions ameliorates Aβo toxicity in primary rat neurons, supporting the idea that inhibiting Tau-SH3 interactions is a potential mechanism by which Tau reduction is protective. We developed several novel tools for studying Tau-SH3 interactions. One is a proximity ligation assay for the most well studied Tau-SH3 interaction, Tau-Fyn. In neurons, endogenous Tau-Fyn interactions were observed primarily along neurites adjacent to Tau-positive neuritic shafts. The second new tool is a cell-permeable peptide inhibitor, Tau-PxxP_5/6_, that competitively inhibits Tau-SH3 interactions in neurons. Blocking Tau-SH3 interactions using this inhibitor reduced Tau-Fyn interactions by PLA and ameliorated multiple measures of Aβ toxicity. The Tau-SH3 interaction inhibitor also reduced Tau phosphorylation by Fyn, without directly reducing Fyn activity. This finding provides further support for the ability of Tau-PxxP_5/6_ to reduce Tau-Fyn interactions, although it is not clear that Fyn phosphorylation of Tau is the mechanism by which Tau-PxxP_5/6_ reduces Aβo toxicity. Our data indicates that Tau-PxxP_5/6_ decreases Fyn phosphorylation of Tau and prevents Aβ toxicity, but we cannot conclude that the former causes the latter. It remains possible that other downstream effects of inhibiting the Tau-Fyn interaction contribute to the prevention of Aβ toxicity by Tau-PxxP_5/6_.

One question that will need to be addressed in ongoing studies is which particular Tau-SH3 interactions are most critical. The field has focused on Fyn as a prototypical SH3-containing binding partner for Tau (35) and considerable evidence suggests a role for Tau-Fyn interactions in AD. First, Tau directly binds Fyn’s SH3 domain domain (23, 24, 36, 37) and both Tau phosphorylation and Tau mutations increase Tau-Fyn interaction (23, 38). Second, Tau and Fyn have converging phenotypes both *in vitro* and *in vivo*.

In primary neurons, genetic knockout of either Tau (25) or Fyn (26) ameliorates Aβo toxicity. *In vivo*, reduction of either Tau (8, 10-14, 39) or Fyn (40) prevents Aβo-induced dysfunction, and Tau reduction decreases susceptibility to network hyperexcitability (11, 18). Third, Fyn overexpression exacerbates Aβo-induced neuronal dysfunction (41) and Tau reduction prevents these Fyn-dependent effects (11). Finally, Tau regulates trafficking of Fyn to the postsynaptic density and preventing this Tau-mediated trafficking of Fyn ameliorates Aβo-induced cognitive deficits, NMDAR activation, and premature mortality in mice (12). Altogether, these findings combined with the results observed here indicate that the Tau-Fyn interaction may have an important role in AD pathogenesis.

Other Tau-SH3 interactions besides Tau-Fyn might also play a role. One recent report suggested that the Tau-Fyn interaction may not be critical for glutamate-induced excitotoxicity (32). This would not be inconsistent with the observations here, particularly because the mechanisms of Aβo- and glutamate-induced toxicity are different. In addition, that study (32) used an overexpression system and manipulated the 7^th^ PxxP binding site, which is likely not the primary Fyn binding site in Tau (23, 36, 37). One group recently completed computational modeling of known Tau-SH3 interactions to determine which of Tau’s PxxP motifs has the highest affinity for each SH3 domain (42). Interestingly, they found that of all 180 PxxP-SH3 combinations they modeled, Tau’s 5^th^ PxxP motif and Fyn’s SH3 domain had the highest binding affinity, which highlights the relevance of targeting Tau’s 5^th^/6^th^ PxxP motifs, as we did with Tau-PxxP_5/6_. It is also important to emphasize that other Tau-SH3 interactions besides Tau-Fyn could mediate the effects we observed. Known Tau-SH3 interactions with clear potential relevance to Aβ-induced abnormalities include Tau’s interaction with the AD genetic risk factor BIN1 (43) and the post-synaptic density protein PSD95 (44). BIN1 regulates propagation of Tau pathology (45) and its *Drosophila* homolog modulates Tau toxicity (46). Tau’s proline-rich domain binds directly to BIN1’s SH3 domain and phosphorylation of Tau regulates this binding (47), highlighting a potential role for the Tau-BIN1 interaction in Tau-mediated AD deficits. A recent report used Tau-BIN1 PLA to identify modulators of Tau-BIN1 interaction (48) and found that phosphorylation of BIN1 modulates the interaction and that there is altered BIN1 phosphorylation in AD brain.

Of note from the *in silico* study of SH3-Tau-PxxP interactions (42) was that the strongest binding for BIN1’s SH3 domain in Tau’s proline-rich domain was the 5^th^ PxxP motif. Additionally, PSD95 interacts via its SH3 domain with Tau at the synapse, forming a complex with Fyn and NMDARs (44), and manipulations that disrupt this Tau/Fyn/PSD95 complex prevent network hyperexcitability and and Aβ toxicity *in vivo* (49). This complex allows Fyn to phosphorylate NMDARs which could lead to increased Ca^2+^ influx and excitotoxicity (50). Thus, multiple different Tau-SH3 interactions could contribute to Aβo-induced neuronal dysfunction.

Another remaining question is whether Aβo promotes increased Tau-SH3 interactions as a mechanism leading to neuronal deficits, or whether Tau-SH3 interactions are permissive of Aβo toxicity without Aβo-induced increases. Either possibility is consistent with the findings here that blocking Tau-SH3 interactions reduces Aβo-induced neuronal dysfunction. On one hand, there is some evidence supporting the idea that Aβo could promote Tau-SH3 interactions. Aβo causes Fyn-mediated local translation of Tau in dendrites (51) and causes Tau translocation into dendritic spines (52), either of which could increase the amount of Tau-SH3 interactions if there are unbound SH3-containing proteins available to bind Tau. However, it is unlikely that there are enough unbound SH3-containing proteins in the dendrites to have Aβ-induced local Tau translation or translocation significantly alter the amount of Tau-SH3 interactions, and it is unclear how this would induce Aβ toxicity.

On the other hand, and perhaps more likely, Tau-SH3 interactions could be permissive of Aβo toxicity, even if Aβo does not lead directly to more or stronger Tau-SH3 interactions. Considering the prototypical SH3-containing protein Fyn, extracellular Aβo activates Fyn when it is anchored to the post-synaptic density so that Fyn phosphorylates its targets. One study suggested that Tau traffics Fyn to dendrites and anchors it to the post-synaptic density (12). Based on this model, inhibiting Tau-SH3 interactions, specifically the Tau-Fyn interaction, could prevent activation of Fyn by Aβo in the postsynapse, preventing signal transduction leading to Aβo toxicity. Going forward, it is important to determine if the Tau-Fyn interaction is indeed the critical Tau-SH3 interaction required for Aβo toxicity and which downstream pathways are activated to cause neuronal dysfunction. Possibilities of key downstream pathways include phosphorylation of Tau or the NMDAR, or activation of the AD risk gene Pyk2, another phosphorylation target of Fyn (53) that is implicated in Aβ-induced dysfunction *in vivo* (54).

Our results support the idea of targeting Tau-SH3 interactions as a novel Tau-directed therapeutic strategy for AD. These data provide preclinical evidence supporting the potential efficacy of inhibiting Tau-SH3 interactions; however, considering the potential of a novel strategy also requires evaluating safety and tractability. While further studies are clearly needed, there is evidence to suggest that inhibiting Tau-SH3 interactions would be safe. Even complete removal of Tau, which de facto would prevent all Tau-SH3 interactions, does not cause dramatic abnormalities in animals (55-59), and even the rather subtle changes in Tau knockout mice are not seen in Tau heterozygous mice with partial Tau reduction (19, 60-62), which better represent the incomplete block that would be provided by Tau-SH3 interaction inhibitors. In fact, Tau reduction with antisense oligonucleotides did not have apparent adverse effects in mouse models (18, 63) and human clinical trials using this approach have been underway since 2017, as yet without reported adverse effects. In terms of tractability, inhibiting protein-protein interactions was once thought to be an unrealistic strategy for small molecule therapies, but advances in drug discovery and design have mitigated these concerns (64). We have already conducted high-throughput screening for Tau-SH3 interaction inhibitors, using a Tau-Fyn interaction assay, and identified multiple series of chemically tractable hits (23). Much is left to learn, but the data presented here provide initial preclinical support for the concept of targeting Tau-SH3 interactions as a therapeutic strategy for AD.

## Conclusions

Tau reduction ameliorates Aβo-induced dysfunction, but there is a need to identify therapeutic strategies that reproduce this effect. Here, we tested the hypothesis that Tau-SH3 interactions contribute to Aβo toxicity and can be targeted therapeutically. We developed a cell-based target engagement assay and a peptide inhibitor that reduces Tau-SH3 interactions. We used PLA to measure endogenous Tau-Fyn interactions throughout the neurites of primary neurons and determined that the peptide inhibitor reduces endogenous Tau-Fyn interactions in neurons. We then used this peptide inhibitor to determine that inhibiting Tau-SH3 interactions ameliorates Aβo toxicity, highlighting the therapeutic potential of inhibiting Tau-SH3 interactions to treat AD.

## List of abbreviations

AD: Alzheimer’s disease.
Aβ: amyloid-β.
Aβo: Aβ oligomer.
SH3: Src-homology 3 domain.
FTD: Frontotemporal dementia.
NMDAR: NMDA receptor.
ASO: antisense oligonucleotide.
NTC: nontargeting control ASO.
MTT: 3-(4,5-dimethylthiazol-2-yl)-2,5-diphenyltetrazolium bromide.
TAT: transactivator of transcription.
PLA: proximity ligation assay.
DIV: days *in vitro*.

## Materials and Methods

### Primary hippocampal neurons

All animal procedures were approved by the Institutional Animal Care and Use Committee of the University of Alabama at Birmingham. Timed-pregnant, albino Sprague-Dawley rats (Charles River) were euthanized by isoflurane anesthesia asphyxiation followed by bilateral thoracotomy. Hippocampal tissue from E19 embryos was harvested on ice in cold Hibernate E media (4°C; Life Technologies A12476-01), and then digested with Papain (20 units/mL, Worthington Biochemical Corporation LK003178) for 10 min at 37°C. After the incubation, neurons were dissociated by manual trituration to a single-cell suspension in Neurobasal medium (ThermoFisher 21103049) supplemented with 1x B-27 (ThermoFisher 17504044), 2 mM L-Glutamine (ThermoFisher 25030081) and 10% premium select fetal bovine serum (Atlanta Biologicals S11550). Neurons were then plated at 30,000 neurons per well (9.0 × 10^4^ neurons per cm^2^) in 200 μL plating medium. Neurons were plated in the inner 60 wells of 96-well plates coated overnight at 4°C with 0.1 mg/mL Poly-D-Lysine (Millipore Sigma P6407) and 0.02 mg/mL laminin (Millipore Sigma L2020) 24-48 hours prior to the neuron harvest, with the outer wells containing autoclaved ultrapure water (MilliQ filtered) to prevent evaporation in wells with neurons. Neurons used for the MTT membrane trafficking assay were plated in clear plastic 96-well plates (Fisher 08-772-2C), and those used for the MAP2 neurite degeneration assay were plated in black plastic, clear bottom 96-well plates (Fisher 07-200-565). Cultures were maintained in a 37°C humidified incubator with 5% CO_2_. 24 hours after plating, 75% media was exchanged for Neurobasal supplemented with B-27 and L-Glutamine but lacking serum. 5μM cytosine β-D-arabinofuranoside (Millipore Sigma C6645) was added at DIV2 to inhibit glial proliferation. 50% media changes were performed weekly with Neurobasal supplemented with B-27 and L-Glutamine until experiments were started at DIV19–21.

### Antisense oligonucleotide application

At DIV14, neurons were treated with either 1 μM Tau ASO (sequence: 5-ATCACTGATTTTGAAGTCCC-3), 1 μM nontargeting control ASO (sequence: 5-CCTTCCCTGAAGGTTCCTCC-3), both ASOs were previously published (18), or 1 μM Fyn ASO (sequence: 5-CACAGCCCATTATCCA-3), which was previously published (29), and ordered from IDT. ASO was left on for one week before performing experiments.

### Inhibitor Peptides

Peptides were synthesized by Peptide 2.0. Tau-PxxP_5/6_, rrrqrrkkrgRSRTPS LPTPPTREPKK, consisted of amino acids 209–225 of Tau, with an N-terminal TAT tag (lower case) for cell permeability. Tau-CTD, rrrqrrkkrgENAKAKTDHG AEIVYKS, consisted of amino acids 380–396 of Tau, also with an N-terminal TAT tag. Peptides were received as lyophilized powder, dissolved in DMSO to 30 mM, then aliquoted and frozen until use, with a final working concentration of 15 μM, unless otherwise stated.

### Western Blotting

For primary neurons, one week after ASO application, neurons were lysed using RIPA buffer (50 mM Tris, pH 7.5, 150 mM NaCl, 5 mM EDTA, 0.1% SDS, 0.1% Triton X100, 0.5% sodium deoxycholate) with protease inhibitors and centrifuged to remove cell debris. HEK-293 cells were lysed with lysis buffer (50 mM Tris, 150 mM NaCl, 5 mM EDTA, 1% Triton X-100, 0.1% sodium deoxycholate) with protease inhibitor and phosphatase inhibitor can centrifuged to remove cell debris. The samples were diluted with LDS buffer (Thermo Fisher NP0007) and sample reducing agent (final concentration 50 mM dithiothreitol, Fisher B0009), and then heated at 70°C for 10 minutes. The samples were then run on 4-12% bis-tris gels (ThermoFisher NW04127BOX) in SDS buffer (25 mMTris base, 190 mM glycine, 0.1% SDS; pH 8.3) with 20 µg of protein per lane. The gels were transferred to Immobilon-FL PVDF membranes (Fisher IPFL00010), and the membranes were blocked for 1 hour in 50% Odyssey blocking buffer (Fisher NC9125955). The membranes were probed with primary antibody overnight at 4C. The next day, the membranes were incubated at room temperature for 1 hour in an IRDye 700 or 800-conjugated secondary antibody (1:20,000 Fisher NC0252291 or Fisher NC0252290) and scanned on an Odyssey Scanner (LI-COR Biotechnology). Bands were quantitated using Image Studio Lite software (LI-COR Biotechnology) and ImageJ. Statistical analysis was performed with GraphPad Prism 8. Antibodies used were: rabbit monoclonal anti-Tau (1:1000, DAKO #A0024), mouse monoclonal anti-NeuN (Millipore MAB377), mouse monoclonal anti-pY18 Tau (1:2500 GeneTex 9G3), rabbit polyclonal anti-SFKpY420 (1:1000, Cell Signalling Technology #2010), rabbit polyclonal anti-SFKpY531 (1:1000, Cell Signalling Technology #2015), mouse monoclonal anti-Fyn (1:500, Fyn15, Santa Cruz sc-434, used for HEK-293 lysate), rabbit polyclonal anti-Fyn (1:1000, Fyn3, Santa Cruz sc-16, used for neuron lysate), mouse monoclonal anti-GAPDH (1:5000, Millipore MAB374).

### Amyloid-β oligomer preparation

Lyophilized recombinant amyloid beta 1-42 (Aβ; California Peptide 641-15) was dissolved in 1,1,1,3,3,3-Hexafluoro-2-propanol (Millipore Sigma 52517), dried overnight, and then stored in a desiccator at -20°C until oligomerization. To prepare oligomers, Aβ was dissolved in fresh, anhydrous DMSO (Millipore Sigma D2650) to 1 mM, vortexed and incubated for 5 minutes at room temperature, and then diluted to 100 μM in 1x PBS on ice. The Aβ solution was then sonicated in an ice bath off and on at 30 sec increments for 15 minutes and then left on ice undisturbed to oligomerize for 24 hours. Immediately prior to use, Aβo was centrifuged at 4°C for 5 minutes at 5,000xg. Vehicle solution was prepared in identical fashion, beginning with 1,1,1,3,3,3-Hexafluoro-2-propanol evaporation, except lacking Aβ peptide.

### Modified MTT assay for membrane trafficking

After 24 hour exposure to Aβo (or vehicle), 100 μM Tetrazoleum salts (MTT; Millipore Sigma M2128) were applied to neurons and incubated at 37°C for 60 minutes. Media from each well was replaced with 15 μL 1xPBS containing 1.6% Tween® 20 (Fisher BP337500) and incubated at room temperature with mild agitation for 5 minutes. The entire sample volume (referred to as “Tween Soluble MTT”) was then transferred to a 384-well white-plastic, clear bottom plate, and 15 μL of isopropanol (Millipore Sigma 190764) was added to the plate. The isopropanol containing plates were incubated at room temperature with mild agitation for 5 minutes, and then (referred to as “Tween Insoluble MTT”) transferred to distinct wells of the same 384-well plate as the associated tween-containing samples. Sample pH was raised by adding 2 μL of 1N NaOH to each well to improve formazan absorbance spectrum. The 384-well plates were spun for 2 minutes at 100xg to eliminate any confounding air bubbles, and then absorbance at 590 nm read on a Synergy2 (BioTek) plate reader with absorbance at 660 nm used as the reference wavelength. The ratio of Tween Soluble MTT to Tween Insoluble MTT was calculated on a per sample basis, and then ratios normalized to the vehicle treated group average per experiment.

### MAP2 immunocytochemistry for neurite retraction assay

After 48 hours exposure to Aβo (or vehicle), neurons were fixed in 1xPBS containing 4% paraformaldehyde and 4% sucrose at 37°C for 45 minutes. Neurons were washed 3×5 mins in 1xPBS, blocked in saponin blocking buffer (5% normal horse serum, 5% normal goat serum, 1% bovine serum albumin, and 0.5% saponin in 1xPBS) for 1 hour, and then incubated overnight at 4°C with MAP2 antibody (ThermoFisher, PA1-10005, 1:5000) in saponin blocking buffer. After overnight incubation, neurons were washed 3×5 mins with saponin rinse buffer (0.5% normal horse serum, 0.5% normal goat serum, 0.05% saponin in 1xPBS). Neurons were washed 4×5minutes with saponin rinse buffer, then secondary antibody (Donkey anti-chicken conjugated with Alexa-594, Jackson Immunoresearch, 702-585-155, 1:2000) applied for 1 hour at room temperature in saponin blocking buffer. Neurons were rinsed 4×5minutes with saponin rinse buffer then washed 5×5mins with 1xPBS. Plates were wrapped in parafilm, protected from light, and stored at 4°C until imaging. Neurons were imaged on the Operetta high-content Imager (PerkinElmer) with a 40x objective collecting 4 adjacent fields per well. Intact neurite length was measured in an unbiased, automated manner using Harmony software (PerkinElmer). Intact neurites have relatively smooth MAP2 labeling, and the neurite detection algorithm excludes neurites that have high contrast along their path (e.g. produced by blebbing). See Supplementary Figure 3 for workflow. For analysis, total intact neurite length from each condition was normalized to vehicle-treated control wells from each plate and compared by Two-Way ANOVA followed by Dunnett’s *post hoc* test with correction for multiple comparisons.

### HEK-293 transfection and inhibitor application

HEK-293 cells were passaged by removing media and washing with 1xPBS, then applying 0.25% trypsin (Fisher 25-200-056) to cells for 3-5 minutes at 37°C to dissociate them from the plate. Trypsin was inactivated by diluting it 1:10 in DMEM (Fisher MT10013CV) supplemented with 10% premium select fetal bovine serum and 1% Pen Strep (ThermoFisher 15140122) and the cells were plated to 10% confluency in 500 μL supplemented DMEM onto 12mm No 1 glass coverslips (Carolina Biological 633029) coated overnight at 4°C with 0.1 mg/mL Poly-D-Lysine and 0.02 mg/mL laminin in a 24-well plate (Fisher 08-772-1). After 24 hours, each well was transfected with 125–500 ng of each plasmid (human Tau tagged with mKate2 and human Fyn tagged with Myc) and 2 uL of Promega Fugene (Fisher) in serum-free DMEM. The following day, 5 μL of 3 mM Tau-PxxP_5/6_ or Tau-CTD (Peptide 2.0) in 10% DMSO in serum-free DMEM was added to each well as appropriate to bring the final concentration to 15 μL of peptide. Twenty-four hours after peptide application, cells were fixed in 1xPBS containing 4% paraformaldehyde and 4% sucrose at room temperature for 30 minutes, then washed 3×5 minutes in 1xPBS and stored at 4°C in the dark until PLA was performed, or lysed for Western blot analysis.

### HEK-293 Proximity Ligation Assay and image analysis

Duolink In Situ Fluoresence kit (Millipore Sigma DUO92014) was used for PLA. Fixed HEK-293 cells on coverslips were permeablized for 10 minutes with 0.25% Triton X-100 (Fisher BP151-500) in 1xPBS at room temperature then blocked in 5% normal goat serum in 1xPBS for 1 hour at room temperature. Coverslips were then incubated overnight at 4°C with Tau5 (1:500) and Fyn3 (Santa Cruz, sc-16, 1:250) antibodies in 1% NGS in 1xPBS in a humidity chamber to prevent evaporation. The next day, coverslips were washed 3×5 minutes in 1xPBS then incubated for 1 hour at 37°C in 8 μL Duolink In Situ PLA Probe Anti-Rabbit PLUS (Millipore Sigma DUO92002) and 8 μL Anti-Mouse MINUS (Millipore Sigma DUO92006) in 24 8 μL 1% NGS in 1xPBS per reaction. Coverslips were washed 2×5 minutes in Duolink In Situ Wash Buffer A at room temperature then incubated for 1 hour at 37°C in 8 μL Duolink In Situ 5x ligation buffer and 1 μL ligase in 31 μL ultrapure DNase/RNase-free distilled water (ThermoFisher 10977015) per reaction. Coverslips were washed 2×2 minutes in Duolink In Situ Wash Buffer A at room temperature then incubated for 100 minutes at 37°C in 8 μL Duolink In Situ 5x amplification buffer and 0.5 uL polymerase in 31.5 μL ultrapure DNase/RNase-free distilled water per reaction. Coverslips were washed 2×10 minutes in Duolink In Situ Wash Buffer B at room temperature then washed for 1 minute in 1% Duolink In Situ Wash Buffer B and for 5 minutes in 1xPBS. To view Fyn, coverslips were incubated for 1 hour at room temperature in secondary antibody (Goat anti-rabbit conjugated with Alexa-647, ThermoFisher, A-21245, 1:1000) in 1% NGS in 1xPBS, then washed 3×5 minutes in 1xPBS. Coverslips were mounted with Duolink In Situ Mounting Medium with DAPI (Sigma) and stored at 4°C while protected from light until imaging. Fluorescent images were taken using an epifluorescent microscope at 60x with four channels: DAPI (nuclei), FITC (PLA), TRITC (Tau-mKate2), and Cy5 (Fyn). Seven images per slide were obtained then analyzed using ImageJ. Because it is challenging to disentangle cells contributing from outside of the field of view, we measured PLA density per field-of-view and took multiple images of each coverslip to accurately measure PLA density for each coverslip. For each of the channels, we subtracted the background and adjusted brightness/contrast equally across treatments. Since Tau filled the cells most evenly, we measured cell area as the area covered by Tau fluorescence by thresholding the TRITC channel. To quantify PLA puncta, we thresholded the FITC channel to view puncta and ran the ImageJ particle analyzer, specifying size and circularity to exclude any non-punctate signal. We then divided the PLA puncta by the area covered by Tau for each image to measure PLA density and normalized to the values for untreated slides from that experiment.

### Primary Neuron PLA

Neurons were grown on coverslips until DIV20, then treated with 15 μM Tau-PxxP_5/6_ or vehicle control for 24 hours and fixed in 4% PFA with 4% glucose in 1x PBS. The same Duolink In Situ Fluoresence kit was used as for the HEK-293 PLA with the only difference being primary and secondary antibodies used. Anti-Tau (1:1000, DAKO #A0024) and anti-Fyn (1:250, Santa Cruz SC-434) primary antibodies were used, and Goat anti-rabbit conjugated with Alexa-594, (ThermoFisher, A-11037, 1:1000) secondary antibody was used to view Tau. Imaging and quantification was completed using the same methods described for the HEK-293 PLA.

### Statistics

Sample sizes were determined based on power calculations to provide 80% power to detect a difference of 20% with 30% standard deviation at an alpha of 0.05 using the effect size from preliminary studies or from literature. All statistical tests were performed using Graphpad Prism 7, and appropriate statistical tests were used based on the number and types of groups for each experiment. Amelioration of Aβo toxicity was defined as a significant interaction between main factors of peptide concentration and Aβo by two-way ANOVA, with a significant difference between vehicle and Aβo at 0 μM peptide and no significant difference between vehicle and Aβo at 15 μM peptide by Dunnet’s *post hoc*.

## Acknowledgements

We thank Yuliya Voskobiynyk for valuable discussions and input throughout the project and for help with data visualization. We also thank the Herskowitz and Arrant labs for valuable discussion and Kelly Chen for assistance with protocol development for immunostaining. Southern Research and the UAB Center for Biophysical Sciences and Engineering graciously provided access to their imaging equipment.

## Declarations

### Availability of supporting data

The data sets used and analyzed during this study are available from the corresponding author upon reasonable request.

### Authors’ contributions

TR, JNC, and EDR conceived of the study. TR, JRR, and JNC developed assays and reagents for the study. TR, JRR, and EDR designed experiments, which were performed by TR, JRR, SJT, and ARA, all of whom completed statistical analysis. Data was analyzed by TR, JRR, and EDR. The manuscript was written by JRR and EDR with input from TR, SJT, and JNC. All authors read and approved the final manuscript.

### Consent for publication

Not applicable.

### Ethics approval

All experiments were conducted in accordance with the guidelines set forth by the Institutional Animal Care and Use Committee at the University of Alabama at Birmingham.

## References

1. Weingarten MD, Lockwood AH, Hwo SY, Kirschner MW. A protein factor essential for microtubule assembly. Proc Natl Acad Sci USA. 1975;72(5):1858–62.

2. Wang Y, Mandelkow E. Tau in physiology and pathology. Nature reviews Neuroscience. 2016;17(1):5–21.

3. Rademakers R, Cruts M, van Broeckhoven C. The role of tau (*MAPT*) in frontotemporal dementia and related tauopathies. Hum Mutat. 2004;24(4):277–95.

4. Hutton M, Lendon CL, Rizzu P, Baker M, Froelich S, Houlden H, et al. Association of missense and 5’-splice-site mutations in tau with the inherited dementia FTDP-17. Nature. 1998;393:702–5.

5. Ihara Y, Nukina N, Miura R, Ogawara M. Phosphorylated tau protein is integrated into paired helical filaments in Alzheimer’s disease. J Biochem. 1986;99(6):1807–10.

6. Grundke-Iqbal I, Iqbal K, Quinlan M, Tung YC, Zaidi MS, Wisniewski HM. Microtubule-associated protein tau. A component of Alzheimer paired helical filaments. J Biol Chem. 1986;261(13):6084–9.

7. Tai XY, Koepp M, Duncan JS, Fox N, Thompson P, Baxendale S, et al. Hyperphosphorylated tau in patients with refractory epilepsy correlates with cognitive decline: a study of temporal lobe resections. Brain. 2016;139(9):2441–55.

8. Roberson ED, Scearce-Levie K, Palop JJ, Yan F, Cheng IH, Wu T, et al. Reducing endogenous tau ameliorates amyloid β-induced deficits in an Alzheimer’s disease mouse model. Science. 2007;316(5825):750–4.

9. Palop JJ, Chin J, Roberson ED, Wang J, Thwin MT, Bien-Ly N, et al. Aberrant excitatory neuronal activity and compensatory remodeling of inhibitory hippocampal circuits in mouse models of Alzheimer’s disease. Neuron. 2007;55(5):697–711.

10. Meilandt WJ, Yu GQ, Chin J, Roberson ED, Palop JJ, Wu T, et al. Enkephalin elevations contribute to neuronal and behavioral impairments in a transgenic mouse model of Alzheimer’s disease. J Neurosci. 2008;28(19):5007–17.

11. Roberson ED, Halabisky B, Yoo JW, Yao J, Chin J, Yan F, et al. Amyloid-β/Fyn–induced synaptic, network, and cognitive impairments depend on tau levels in multiple mouse models of Alzheimer’s disease. J Neurosci. 2011;31(2):700–11.

12. Ittner LM, Ke YD, Delerue F, Bi M, Gladbach A, van Eersel J, et al. Dendritic function of tau mediates amyloid-beta toxicity in Alzheimer’s disease mouse models. Cell. 2010;142(3):387–97.

13. Nussbaum JM, Schilling S, Cynis H, Silva A, Swanson E, Wangsanut T, et al. Prion-like behaviour and tau-dependent cytotoxicity of pyroglutamylated amyloid-β. Nature. 2012;485(7400):651–5.

14. Leroy K, Ando K, Laporte V, Dedecker R, Suain V, Authelet M, et al. Lack of tau proteins rescues neuronal cell death and decreases amyloidogenic processing of APP in APP/PS1 mice. Am J Pathol. 2012;181(6):1928–40.

15. Holth JK, Bomben VC, Reed JG, Inoue T, Younkin L, Younkin SG, et al. Tau loss attenuates neuronal network hyperexcitability in mouse and Drosophila genetic models of epilepsy. J Neurosci. 2013;33(4):1651–9.

16. Gheyara AL, Ponnusamy R, Djukic B, Craft RJ, Ho K, Guo W, et al. Tau reduction prevents disease in a mouse model of Dravet syndrome. Ann Neurol. 2014;76(3):443–56.

17. Palop JJ, Mucke L. Epilepsy and cognitive impairments in Alzheimer disease. Arch Neurol. 2009;66(4):435–40.

18. Devos SL, Goncharoff DK, Chen G, Kebodeaux CS, Yamada K, Stewart FR, et al. Antisense reduction of tau in adult mice protects against seizures. J Neurosci. 2013;33(31):12887–97.

19. Li Z, Hall AM, Kelinske M, Roberson ED. Seizure resistance without parkinsonism in aged mice after tau reduction. Neurobiol Aging. 2014;35(11):2617–24.

20. Congdon EE, Sigurdsson EM. Tau-targeting therapies for Alzheimer disease. Nature reviews Neurology. 2018;14(7):399–415.

21. Cummings J, Blennow K, Johnson K, Keeley M, Bateman RJ, Molinuevo JL, et al. Anti-Tau Trials for Alzheimer’s Disease: A Report from the EU/US/CTAD Task Force. J Prev Alzheimers Dis. 2019;6(3):157–63.

22. Gong CX, Liu F, Grundke-Iqbal I, Iqbal K. Post-translational modifications of tau protein in Alzheimer’s disease. J Neural Transm. 2005;112(6):813–38.

23. Cochran JN, Diggs PV, Nebane NM, Rasmussen L, White EL, Bostwick R, et al. AlphaScreen HTS and live-cell bioluminescence resonance energy transfer (BRET) assays for identification of Tau-Fyn SH3 interaction inhibitors for Alzheimer disease. J Biomol Screen. 2014;19(10):1338–49.

24. Lee G, Newman ST, Gard DL, Band H, Panchamoorthy G. Tau interacts with src-family non-receptor tyrosine kinases. J Cell Sci. 1998;111:3167–77.

25. Rapoport M, Dawson HN, Binder LI, Vitek MP, Ferreira A. Tau is essential to β-amyloid-induced neurotoxicity. Proc Natl Acad Sci USA. 2002;99(9):6364–9.

26. Lambert MP, Barlow AK, Chromy BA, Edwards C, Freed R, Liosatos M, et al. Diffusible, nonfibrillar ligands derived from Aβ1-42 are potent central nervous system neurotoxins. Proc Natl Acad Sci USA. 1998;95:6448–53.

27. Kojima N, Ishibashi H, Obata K, Kandel ER. Higher seizure susceptibility and enhanced tyrosine phosphorylation of N-methyl-D-aspartate receptor subunit 2B in fyn transgenic mice. Learn Mem. 1998;5(6):429–45.

28. Ittner A, Ittner LM. Dendritic Tau in Alzheimer’s Disease. Neuron. 2018;99(1):13–27.

29. Girasol A, Albuquerque GG, Mansour E, Araújo EP, Degasperi G, Denis RG, et al. Fyn Mediates Leptin Actions in the Thymus of Rodents. PLOS ONE. 2009;4(11):e7707.

30. Liu Y, Schubert D. Cytotoxic amyloid peptides inhibit cellular 3-(4,5-dimethylthiazol-2-yl)-2,5-diphenyltetrazolium bromide (MTT) reduction by enhancing MTT formazan exocytosis. J Neurochem. 1997;69(6):2285–93.

31. Izzo NJ, Staniszewski A, To L, Fa M, Teich AF, Saeed F, et al. Alzheimer’s therapeutics targeting amyloid beta 1-42 oligomers I: Abeta 42 oligomer binding to specific neuronal receptors is displaced by drug candidates that improve cognitive deficits. PloS one. 2014;9(11):e111898–e.

32. Miyamoto T, Stein L, Thomas R, Djukic B, Taneja P, Knox J, et al. Phosphorylation of tau at Y18, but not tau-fyn binding, is required for tau to modulate NMDA receptor-dependent excitotoxicity in primary neuronal culture. Mol Neurodegener. 2017;12(1):41.

33. Knox R, Jiang X. Fyn in Neurodevelopment and Ischemic Brain Injury. Dev Neurosci. 2015;37(4-5):311–20.

34. Wang J, Carnicella S, Phamluong K, Jeanblanc J, Ronesi JA, Chaudhri N, et al. Ethanol induces longterm facilitation of NR2B-NMDA receptor activity in the dorsal striatum: implications for alcohol drinking behavior. The Journal of neuroscience : the official journal of the Society for Neuroscience. 2007;27(13):3593–602.

35. Lee G. Tau and src family tyrosine kinases. Biochim Biophys Acta. 2005;1739(2–3):323–30.

36. Usardi A, Pooler AM, Seereeram A, Reynolds CH, Derkinderen P, Anderton B, et al. Tyrosine phosphorylation of tau regulates its interactions with Fyn SH2 domains, but not SH3 domains, altering the cellular localization of tau. FEBS J. 2011;278(16):2927–37.

37. Lau DH, Hogseth M, Phillips EC, O’Neill MJ, Pooler AM, Noble W, et al. Critical residues involved in tau binding to fyn: implications for tau phosphorylation in Alzheimer’s disease. Acta Neuropathol Commun. 2016;4(1):49.

38. Bhaskar K, Yen SH, Lee G. Disease-related modifications in tau affect the interaction between Fyn and Tau. J Biol Chem. 2005;280(42):35119–25.

39. Palop JJ, Chin J, Roberson ED, Wang J, Thwin MT, Bien-Ly N, et al. Aberrant excitatory neuronal activity and compensatory remodeling of inhibitory hippocampal circuits in mouse models of Alzheimer’s disease. Neuron. 2007;55:697–711.

40. Chin J, Palop JJ, Yu GQ, Kojima N, Masliah E, Mucke L. Fyn kinase modulates synaptotoxicity, but not aberrant sprouting, in human amyloid precursor protein transgenic mice. J Neurosci. 2004;24(19):4692–7.

41. Chin J, Palop JJ, Puolivali J, Massaro C, Bien-Ly N, Gerstein H, et al. Fyn kinase induces synaptic and cognitive impairments in a transgenic mouse model of Alzheimer’s disease. J Neurosci. 2005;25(42):9694–703.

42. Wang X, Chang C, Wang D, Hong S. Systematic profiling of SH3-mediated Tau-Partner interaction network in Alzheimer’s disease by integrating in silico analysis and in vitro assay. Journal of molecular graphics & modelling. 2019;90:265–72.

43. Lambert JC, Ibrahim-Verbaas CA, Harold D, Naj AC, Sims R, Bellenguez C, et al. Meta-analysis of 74,046 individuals identifies 11 new susceptibility loci for Alzheimer’s disease. Nat Genet. 2013;45(12):1452–8.

44. Mondragón-Rodríguez S, Trillaud-Doppia E, Dudilot A, Bourgeois C, Lauzon M, Leclerc N, et al. Interaction of endogenous tau protein with synaptic proteins is regulated by *N*-Methyl-D-aspartate receptor-dependent tau phosphorylation. J Biol Chem. 2012;287(38):32040–53.

45. Calafate S, Flavin W, Verstreken P, Moechars D. Loss of Bin1 Promotes the Propagation of Tau Pathology. Cell reports. 2016;17(4):931–40.

46. Chapuis J, Hansmannel F, Gistelinck M, Mounier A, Van Cauwenberghe C, Kolen KV, et al. Increased expression of BIN1 mediates Alzheimer genetic risk by modulating tau pathology. Mol Psychiatry. 2013;18(11):1225–34.

47. Sottejeau Y, Bretteville A, Cantrelle FX, Malmanche N, Demiaute F, Mendes T, et al. Tau phosphorylation regulates the interaction between BIN1’s SH3 domain and Tau’s proline-rich domain. Acta neuropathologica communications. 2015;3:58.

48. Sartori M, Mendes T, Desai S, Lasorsa A, Herledan A, Malmanche N, et al. BIN1 recovers tauopathy-induced long-term memory deficits in mice and interacts with Tau through Thr(348) phosphorylation. Acta Neuropathol. 2019.

49. Ittner A, Chua SW, Bertz J, Volkerling A, van der Hoven J, Gladbach A, et al. Site-specific phosphorylation of tau inhibits amyloid-beta toxicity in Alzheimer’s mice. Science. 2016;354(6314):904–8.

50. Liu J, Chang L, Song Y, Li H, Wu Y. The Role of NMDA Receptors in Alzheimer’s Disease. Frontiers in Neuroscience. 2019;13(43).

51. Li C, Gotz J. Somatodendritic accumulation of Tau in Alzheimer’s disease is promoted by Fyn-mediated local protein translation. Embo j. 2017;36(21):3120–38.

52. Frandemiche ML, De Seranno S, Rush T, Borel E, Elie A, Arnal I, et al. Activity-dependent tau protein translocation to excitatory synapse is disrupted by exposure to amyloid-Beta oligomers. J Neurosci. 2014;34(17):6084–97.

53. Qian J, Colmers WF, Saggau P. Inhibition of synaptic transmission by neuropeptide Y in rat hippocampal area CA1: Modulation of presynaptic Ca2+ entry. J Neurosci. 1997;17(21):8169–77.

54. Salazar SV, Cox TO, Lee S, Brody AH, Chyung AS, Haas LT, et al. Alzheimer’s Disease Risk Factor Pyk2 Mediates Amyloid-beta-Induced Synaptic Dysfunction and Loss. J Neurosci. 2019;39(4):758–72.

55. Harada A, Oguchi K, Okabe S, Kuno J, Terada S, Ohshima T, et al. Altered microtubule organization in small-calibre axons of mice lacking *tau* protein. Nature. 1994;369(6480):488–91.

56. Dawson HN, Ferreira A, Eyster MV, Ghoshal N, Binder LI, Vitek MP. Inhibition of neuronal maturation in primary hippocampal neurons from tau deficient mice. J Cell Sci. 2001;114:1179–87.

57. Tucker KL, Meyer M, Barde YA. Neurotrophins are required for nerve growth during development. Nat Neurosci. 2001;4(1):29–37.

58. Fujio K, Sato M, Uemura T, Sato T, Sato-Harada R, Harada A. 14-3-3 proteins and protein phosphatases are not reduced in tau-deficient mice. Neuroreport. 2007;18(10):1049–52.

59. van Hummel A, Bi M, Ippati S, van der Hoven J, Volkerling A, Lee WS, et al. No Overt Deficits in Aged Tau-Deficient C57Bl/6.Mapttm1(EGFP)Kit GFP Knockin Mice. PLOS ONE. 2016;11(10):e0163236.

60. Ikegami S, Harada A, Hirokawa N. Muscle weakness, hyperactivity, and impairment in fear conditioning in tau-deficient mice. Neurosci Lett. 2000;279(3):129–32.

61. Lei P, Ayton S, Moon S, Zhang Q, Volitakis I, Finkelstein DI, et al. Motor and cognitive deficits in aged tau knockout mice in two background strains. Molecular neurodegeneration. 2014;9:29.

62. Morris M, Hamto P, Adame A, Devidze N, Masliah E, Mucke L. Age-appropriate cognition and subtle dopamine-independent motor deficits in aged *Tau* knockout mice. Neurobiol Aging. 2013;34(6):1523–9.

63. DeVos SL, Miller RL, Schoch KM, Holmes BB, Kebodeaux CS, Wegener AJ, et al. Tau reduction prevents neuronal loss and reverses pathological tau deposition and seeding in mice with tauopathy. Science translational medicine. 2017;9(374):eaag0481.

64. Arkin MR, Tang Y, Wells JA. Small-molecule inhibitors of protein-protein interactions: progressing toward the reality. Chem Biol. 2014;21(9):1102–14.

