## Supplementary Figures for "A peptide inhibitor of Tau-SH3 interactions ameliorates amyloid-β toxicity"

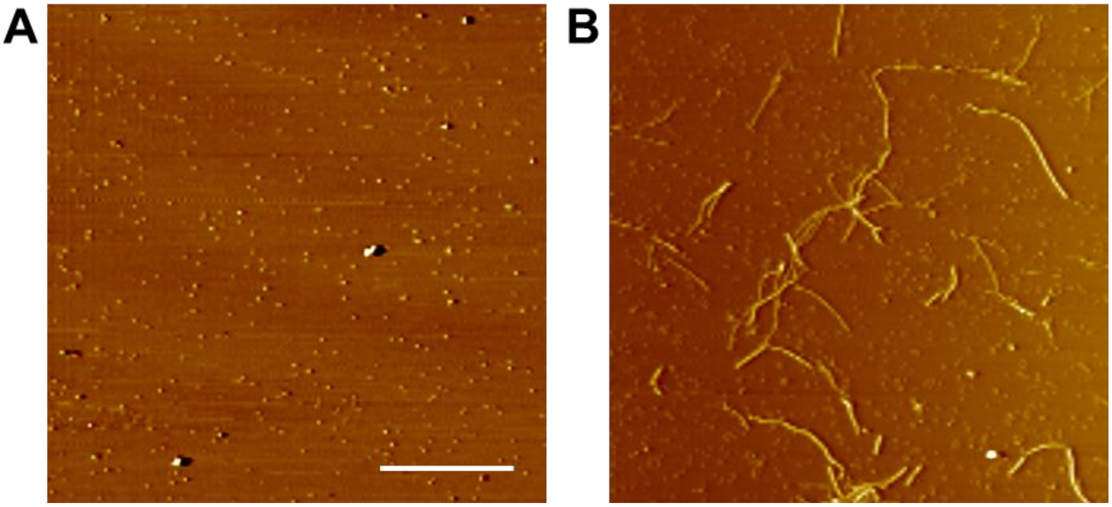


**Supplementary Figure 1: Amyloid-β oligomer and fibril characterization.**

**A:** Representative atomic force micrographs of synthetic amyloid-$\beta_{1-42}$ oligomers. **B:** Representative atomic force micrographs of synthetic amyloid-$\beta_{1-42}$ fibrils after incubation for 48 hours. Aβo and fibrils were produced as procedure detailed in the materials and methods. Scale bar = 1 µm.

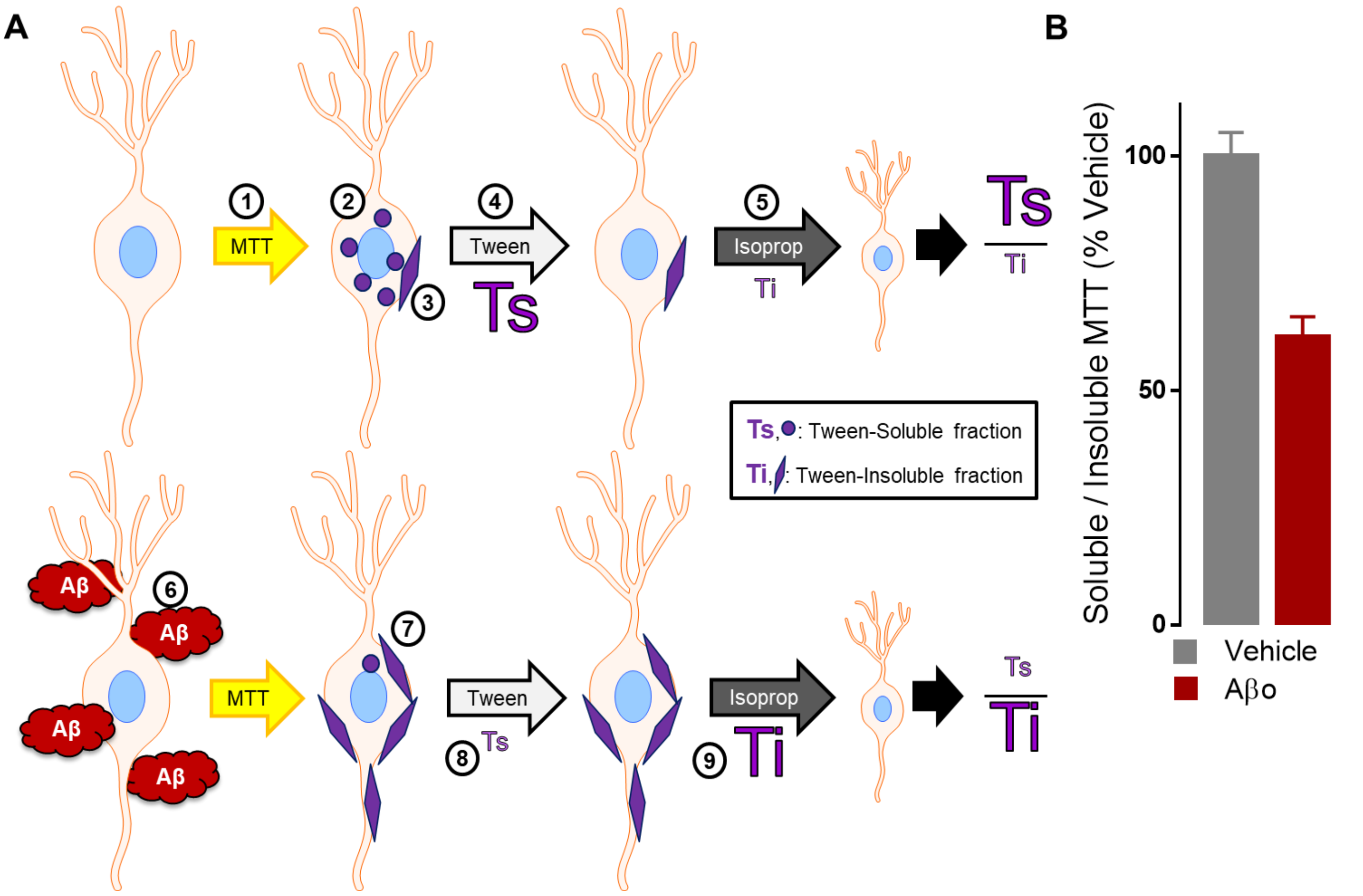


**Supplementary Figure 2: Modified MTT assay workflow.**

Neurons were grown in a 96-well plate to provide in-plate controls and testing every condition on each plate. **A:** Neurons are treated with MTT, which forms a yellow solution (1). Neurons uptake the MTT and metabolize it into formazan, which is purple. This formazan is initially in a soluble form in intracellular vesicles (2). Some of these vesicles are exocytosed, releasing the formazan, where it forms insoluble crystals on the cell membrane of the neuron (3). To measure the amount of soluble intracellular formazan, the detergent Tween-20 is applied (4), dissolving any vesicular formazan. The Tween is removed, representing the Tween-soluble MTT fraction (Ts, indicated by intracellular purple circles). After removing Tween, isopropanol is applied, which dissolves any crystalized formazan (5). The isopropanol is then removed, representing the Tween-insoluble fraction (Ti, indicated by extracellular purple crystals). Pretreatment with Aβo (6) causes the neuron to exocytose more of the formazan, leading to more crystal formation (7). This leads to a smaller Ts (8) and a larger Ti (9) after Aβo application. The amount of formazan in each fraction can be measured using a plate reader. **B:** The ratio of Tween-soluble to Tween-insoluble fractions is a readout of Aβo-induced membrane trafficking abnormalities (same data as in Figure 5).

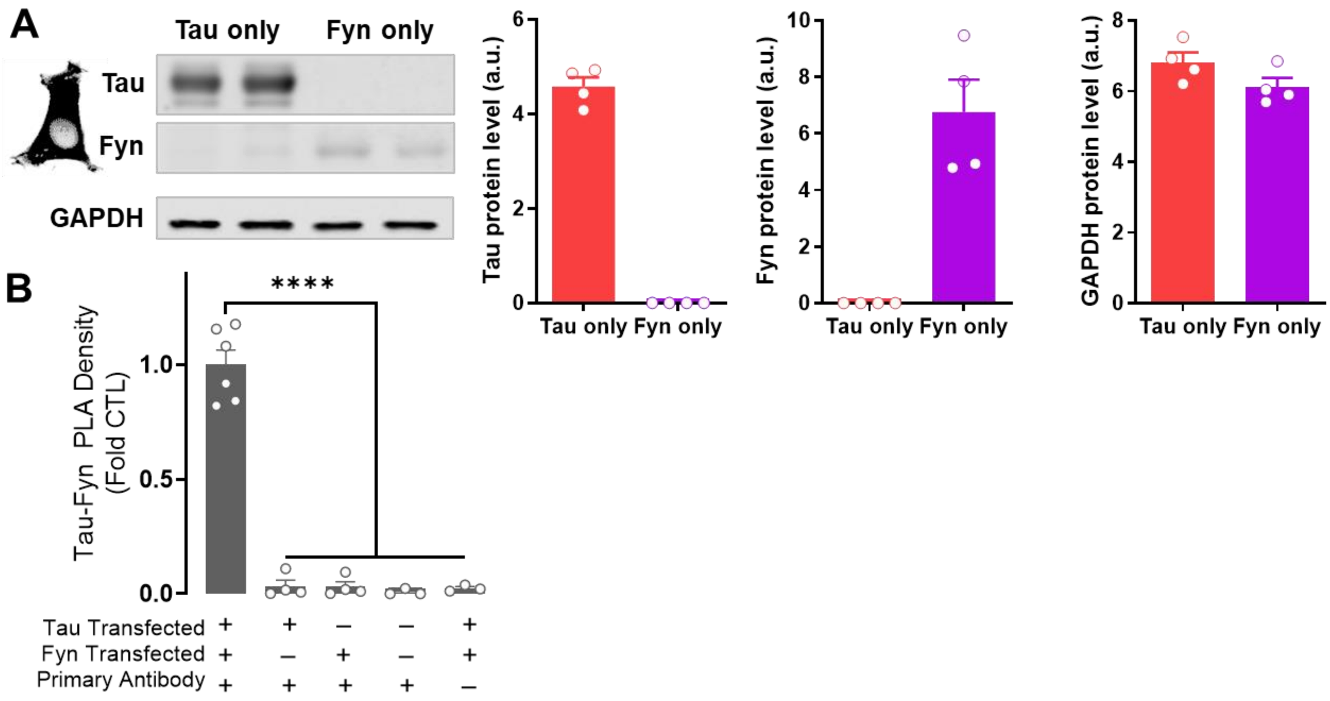


**Supplementary Figure 3: Both Tau and Fyn are required for Tau-Fyn PLA signal in HEK-293 cells.**

HEK-293 cells were transfected with Tau, Fyn or both Tau and Fyn. **A:** Western blot analysis demonstrates that cells transfected with Tau only had increased Tau (Student's t-test, $t(6) = 23.20$, $p < 0.0001$) and cells transfected with Fyn only had increased Fyn (Student's t-test, $t(5) = 5.91$, $p = 0.001$), without no difference in GAPDH levels between groups (Student's t-test, $t(6) = 1.85$, $p = 0.11$). **B:** Tau-Fyn PLA signal was only seen in HEK-293 cells transfected with both Tau and Fyn with primary antibodies applied, while there was only very little background puncta-like fluorescence in cells transfected with only Tau, only Fyn, neither, or transfected with both but without primary antibody, verifying that the Tau-Fyn PLA signal is measuring Tau-Fyn interaction (ANOVA, $F = 71.4$, $p < 0.0001$, all groups different from +/+/+ group by Dunnett's multiple comparisons $p < 0.0001$).

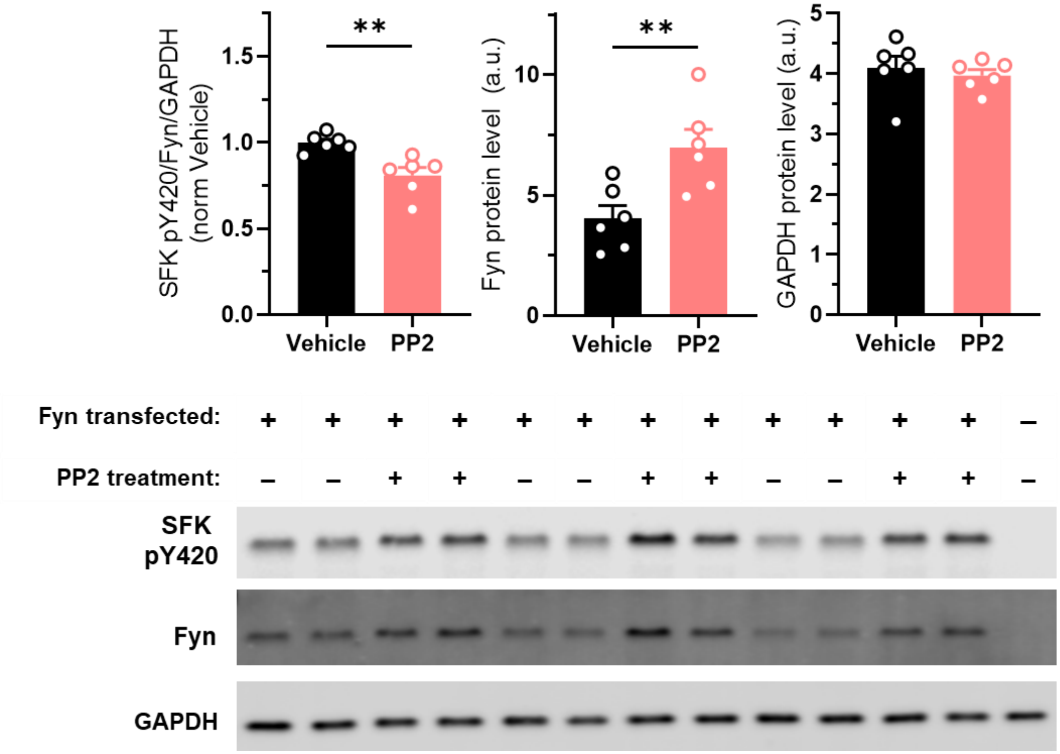


**Supplementary Figure 4: PP2 reduces Fyn kinase activity.**

HEK-293 cells transfected with Fyn were treated with 5 uM PP2 for 3 hours before lysing for Western Blot analysis. PP2 increased Fyn expression (Student's t-test, $t(10) = 3.21$, $p = 0.009$), likely due to compensation for lost Fyn activation, but decreased Fyn kinase activation measured as SFK pY420 (Student's t-test, $t(10) = 3.78$, $p = 0.0036$) without affecting GAPDH levels Student's t-test, $t(10) = 0.57$, $p = 0.58$).

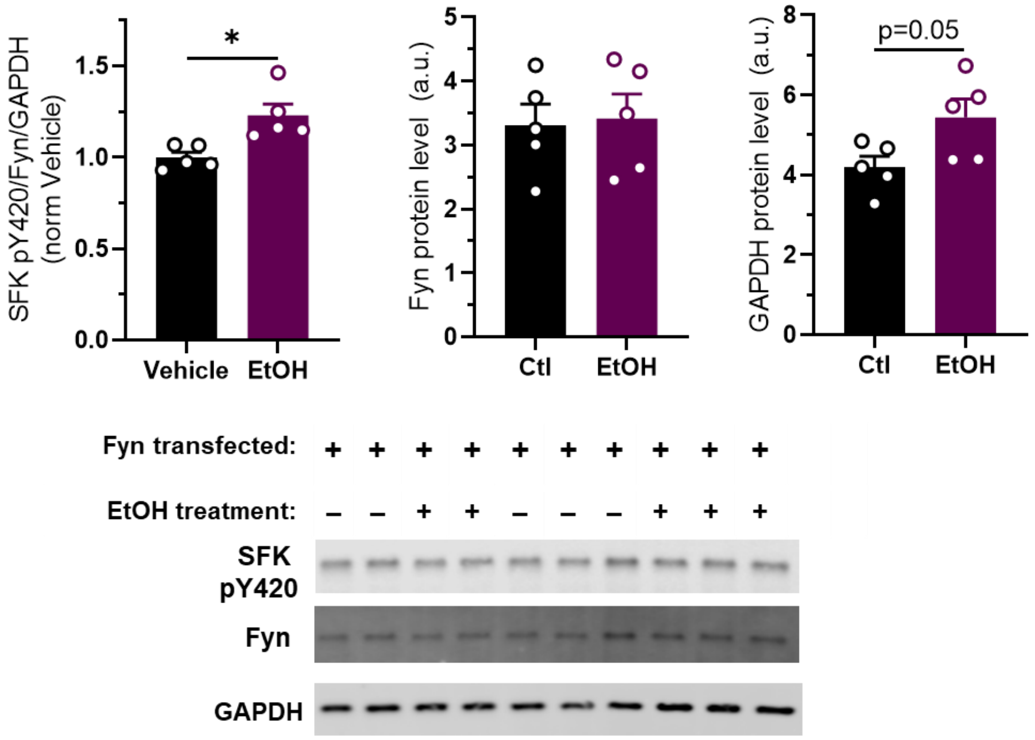


**Supplementary Figure 5: Ethanol increases Fyn kinase activity.**

HEK-293 cells transfected with Fyn were treated with 10 mM EtOH for 30 minutes before lysing for Western Blot analysis. EtOH increased Fyn kinase activation measured as SFK pY420 (Student's t-test, $t(8) = 3.35$, $p = 0.010$) without affecting Fyn expression (Student's t-test, $t(8) = 0.22$, $p = 0.83$). GAPDH protein level was slightly increased in the EtOH group, likely due to loading (Student's t-test, $t(8) = 2.32$, $p = 0.05$), though this was barely a significant change, and, importantly, SFK pY420 was normalized to GAPDH.

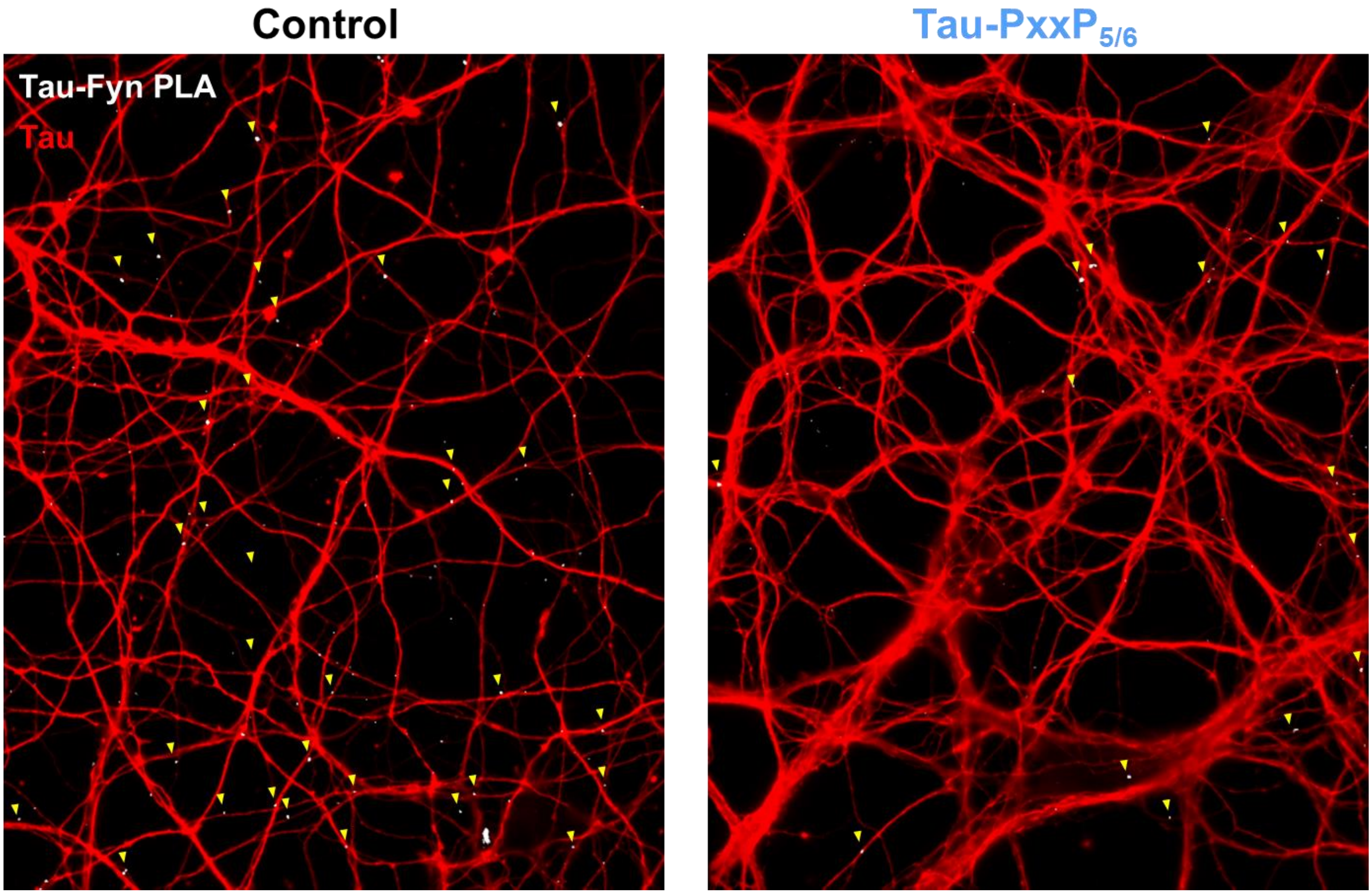


**Supplementary Figure 6: Lower magnification images of Tau-Fyn PLA in primary neurons treated with Tau-PxxP$_{5/6}$.**

Lower magnification images of neurons demonstrating that Tau-PxxP$_{5/6}$ does not adversely affect neurons and decreases Tau-Fyn interaction.

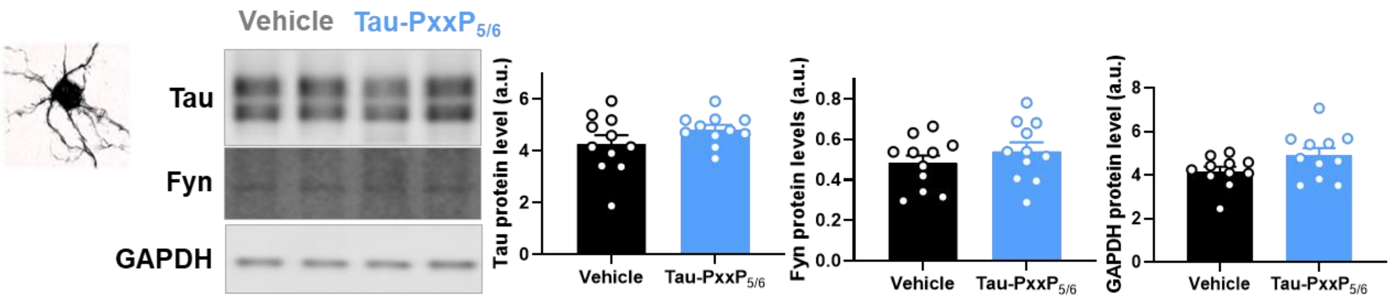


**Supplementary Figure 7: Tau-PxxP$_{5/6}$ does not affect Tau or Fyn levels in primary neurons.**

DIV 20 primary neurons were treated with 15 μM Tau-PxxP$_{5/6}$ for 24 hours then lysed for Western blot analysis. Tau-PxxP$_{5/6}$ did not change levels of Tau (Student's t-test, t(20) = 1.46, $p$ = 0.16), Fyn (Student's t-test, t(20) = 1.00, $p$ = 0.33), or GAPDH (Student's t-test, t(20) = 1.91, $p$ = 0.07).

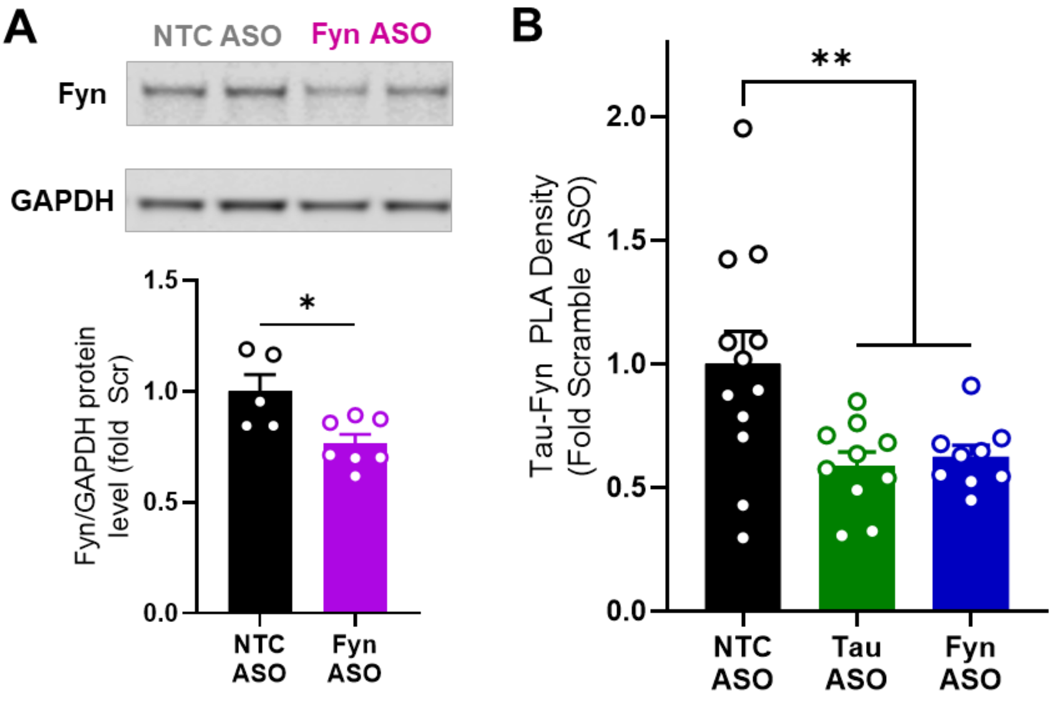


**Supplementary Figure 8: Treatment with Tau or Fyn antisense oligonucleotides reduces Tau-Fyn PLA in primary neurons.**

**A:** Treating neurons with 1 μM Fyn ASO reduced Fyn levels by 24% (Student's t-test, t(10) = 2.96, $p = 0.014$). **B:** DIV 14 primary neurons were treated with 1 μM nontargeting control (NTC), Tau, or Fyn ASOs for 1 week, then fixed for PLA. Reducing either Tau or Fyn reduced Tau-Fyn PLA signal (ANOVA, $F(2,28) = 5.96$, $p = 0.007$, Dunnett's multiple comparisons Scr vs. Tau $p = 0.008$ and Scr vs. Fyn $p = 0.02$).

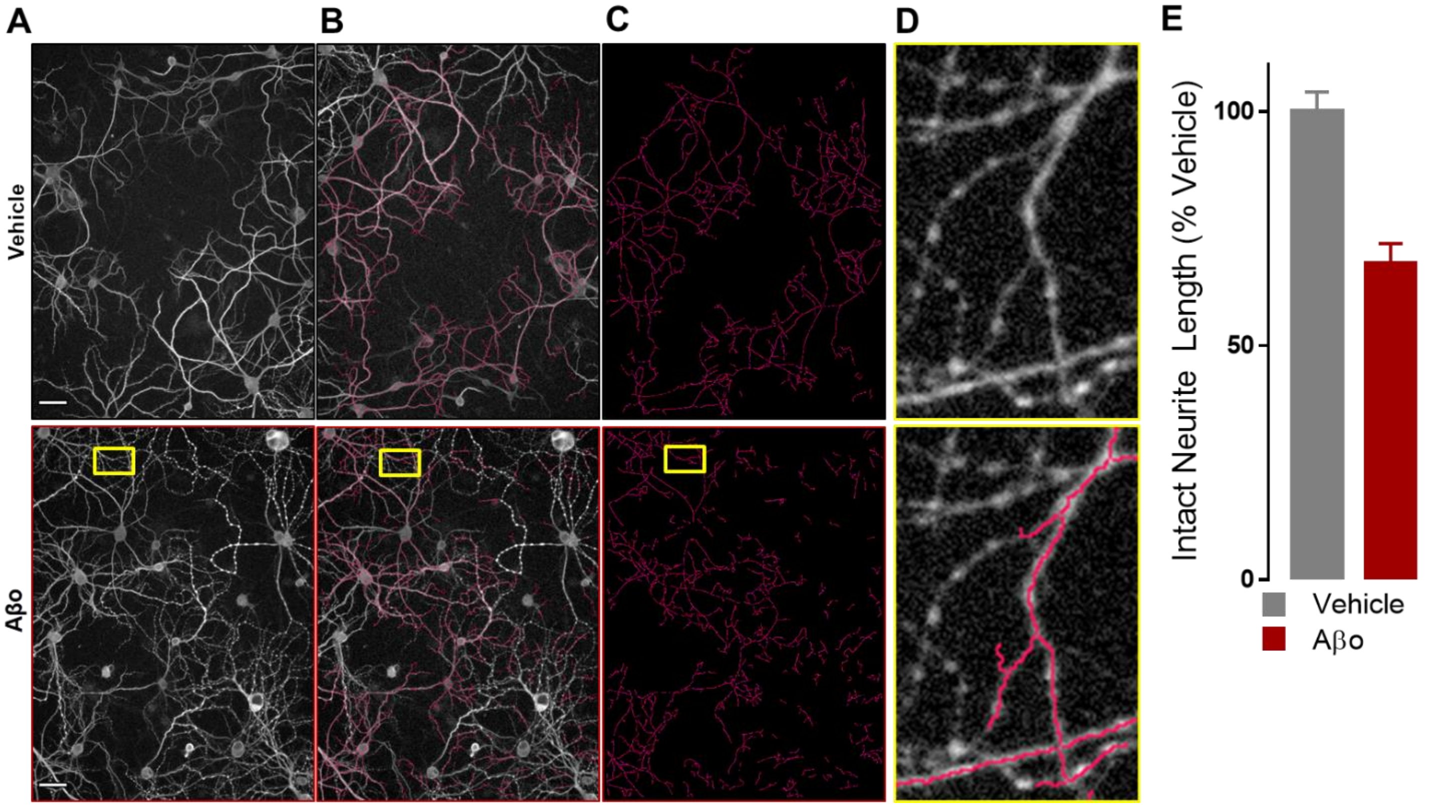


**Supplementary Figure 9: High-content MAP2 imaging and unbiased quantification assay.**

Neurons were grown in a 96-well plate to provide in-plate controls and testing each condition for each plate. After neurons were fixed and immunofluorescently stained for MAP2, four images were collected from each well using an Operetta high-content imager. After images were collected, intact neurite length was measured with an unbiased algorithm using Harmony software. First, Harmony identifies cell bodies based on thresholding, area, and contrast (Find Cells Method B). After cell bodies are identified, the algorithm uses staining morphology to measure intact neurite length in each image (CSIRO Neurite Analysis 2.0). Thus, only intact, unblebbed neurites are measured. **A:** MAP2 staining of neurons. Aβo causes blebbing. Scale bar = 100 μm. **B***:* Neurite measurement superimposed on MAP2 staining. **C**: Neurite measurements. **D**: Inset of images of Aβo-treated neurites on left showing measurement of intact neurites but not blebbed neurites (insets were rotated 90° clockwise). **E:** Aβo decreases intact neurite length (same data as in Figure 6).
